# A comprehensive multi-omics signature of doxorubicin-induced cellular senescence in the postmenopausal human ovary

**DOI:** 10.1101/2024.10.02.616143

**Authors:** Pooja Raj Devrukhkar, Mark A. Watson, Bikem Soygur, Fei Wu, Joanna Bons, Hannah Anvari, Tomiris Atazhanova, Nicolas Martin, Tommy Tran, Kevin Schneider, Jacob P. Rose, Elisheva Shanes, Mary Ellen G. Pavone, Judith Campisi, David Furman, Simon Melov, Birgit Schilling, Francesca E. Duncan

**Author notes:** Co-Correspondence: Francesca E. Duncan, Ph.D. Department of Obstetrics and Gynecology, Feinberg School of Medicine, Northwestern University, Chicago, IL, 60611 USA Birgit Schilling, Ph.D. Buck Institute for Research on Aging, Novato, CA 94945, USA.

## Abstract

A major aging hallmark is the accumulation of cellular senescence burden. Over time senescent cells contribute to tissue deterioration through chronic inflammation and fibrosis driven by the Senescence-Associated Secretory Phenotype (SASP). The human ovary is one of the first organs to age, and prominent age-related fibroinflammation within the ovarian microenvironment is consistent with the presence of senescent cells, but these cells have not been characterized in the human ovary. We thus established a doxorubicin-induced model of cellular senescence to establish a “senotype” (gene/protein signature of a senescence cell state) for ovarian senescent cells. Explants of human postmenopausal ovarian cortex and medulla were treated with doxorubicin for 24 hours followed by culture for up to 10 days in a doxorubicin-free medium. Tissue viability was confirmed by histology, lack of apoptosis, and continued glucose consumption by explants. Single nuclei sequencing and proteomics revealed an unbiased signature of ovarian senescence. We identified distinct senescence profiles for the cortex and medulla, driven predominantly by epithelial and stromal cells. Proteomics uncovered subregional differences in addition to 120 proteins common to the cortex and medulla SASP. Integration of transcriptomic and proteomic analyses revealed 26 shared markers, defining a senotype of doxorubicin-induced senescence unique to the postmenopausal ovary. A subset of these proteins: Lumican, SOD2, MYH9, and Periostin were mapped onto native tissue to reveal compartment-specific localization. This senotype will help determine the role of cellular senescence in ovarian aging, inform biomarker development to identify, and therapeutic applications to slow or reverse ovarian aging, senescence, and cancer.

## Introduction

The ovary is one of the first organs to age in humans (Broekmans et al., 2009). Ovarian aging is associated with a decline in oocyte number and quality, along with increased stromal fibroinflammation, all of which can contribute to infertility (Amargant et al., 2020; Briley et al., 2016; Duncan et al., 2018; Foley et al., 2021; Isola et al., 2024; Landry et al., 2022; Lliberos et al., 2021; Machlin et al., 2021; McCloskey et al., 2020; Umehara et al., 2022). In fact, as more women are delaying childbearing, the reproductive complications associated with advanced age are tangible and often require the use of medically assisted reproduction (Seshadri et al., 2021; Tierney & Guzzo, 2023). Additionally, the decline in gonadal hormones that occurs with peri-menopause and menopause is associated with an increased risk of morbidities such as osteoporosis, cardiovascular diseases, and cognitive dysfunction, thus significantly affecting general health and quality of life (Gracia & Freeman, 2018; Monteleone et al., 2018). Even though improvements in healthcare have increased the average lifespan, the age of menopause has remained relatively constant. As a result, more women are living longer in a post-menopausal state and experiencing the negative sequelae of reduced endocrine function (Monteleone et al., 2018; Pinheiro et al., 2019). Thus, there is a need to better understand the mechanisms underlying ovarian aging to inform therapeutic interventions to promote ovarian longevity and overall health.

Cellular senescence is a fundamental mechanism of mammalian aging characterized by proliferative arrest in response to DNA damage, oxidative stress, and genotoxic insults (Di Micco et al., 2021; Karabicici et al., 2021; López-Otín et al., 2023). While in proliferative arrest, senescent cells maintain metabolic activity and produce a pro-inflammatory secretome known as the senescence-associated secretory phenotype (SASP) consisting of chemokines, interleukins, proteases, and other factors (Di Micco et al., 2021; Kumari & Jat, 2021; Wiley & Campisi, 2021). The SASP confers in senescent cells the ability to locally and distally affect tissue microenvironments, alter tissue homeostasis, and cause tissue damage (Di Micco et al., 2021; Wiley & Campisi, 2021). Throughout life, exposure to damaging stimuli increases the production and accumulation of senescent cells, resulting in chronic inflammation and fibrosis and cumulatively contributing to aging and age-related pathologies (Childs et al., 2015; Mylonas & O’Loghlen, 2022).

Currently, there is no universal marker for identifying senescent cells, and their characteristics vary by the inducer, cell type, and tissue and culture microenvironment (Hernandez-Segura et al., 2018; Herranz & Gil, 2018; Suryadevara et al., 2024). Furthermore, the SASP is highly complex and plastic, varying in composition depending on the cell type and inducer (N. Basisty et al., 2020; Coppé et al., 2010). Although bona fide senescent cells have not been systematically identified or characterized in the mammalian ovary, there is accumulating evidence that senescent cells likely exist. For example, two highly conserved proteins involved in cellular senescence, p21^CIP1^ and p16^INK4a,^ exhibit increased expression with age in human and mouse ovaries (Ansere et al., 2021; Krishnamurthy et al., 2004). The mammalian ovarian microenvironment, consisting of the stroma and follicular fluid, also assumes a prominent fibrotic and inflammatory phenotype with advanced reproductive age, consistent with fibroinflammation caused by senescent cells in other organs (Amargant et al., 2020; Isola et al., 2024; Landry et al., 2022; Lliberos et al., 2021; Machlin et al., 2021; McCloskey et al., 2020; Mylonas & O’Loghlen, 2022; Umehara et al., 2022). Additionally, mitochondrial dysfunction, a hallmark of aging and a potential driver of cellular senescence has been identified as a key contributor to fibrosis and inflammation-induced ovarian decline, particularly in ovarian stroma (Hernandez-Segura et al., 2018; Korolchuk et al., 2017; López-Otín et al., 2023; Umehara et al., 2022; Wiley et al., 2016). Taken together, these findings are highly suggestive of an increasing burden of senescent cells contributing to ovarian aging.

The current gold standard is to use a combination of markers to identify a senescent state (Suryadevara et al., 2024), underscoring the need for cell, tissue, and stimuli-specific markers of cellular senescence. Therefore, the goal of this study was to use a doxorubicin-induced model to establish a cellular senescence senotype in the post-menopausal human ovary that could then be mapped onto native tissue (Figure 1a). Doxorubicin, a DNA-damaging drug, causes follicular apoptosis, microvascular damage, and stromal cell necrosis when used at higher doses typically used in chemotherapeutic regimens (Ben-Aharon et al., 2010; F. Li et al., 2014; Morgan et al., 2013; Soleimani et al., 2011). However, at lower doses, it induces cellular senescence in fibroblasts, cardiomyocytes, and skin (Alimirah et al., 2020; Altieri et al., 2016; Kitada et al., 2019; Marques et al., 2020). We developed an explant culture model of post-menopausal human ovarian tissue that allowed us to interrogate cellular senescence in the context of a complex, intact tissue with cellular heterogeneity. Ovarian explants were treated with or without 24-hour doxorubicin followed by culture in a doxorubicin-free medium for up to 10 days (Figure 1a). Tissue viability was confirmed by histology, cell death, and glucose consumption. Senescence induction was assessed through SA-βGal staining as well as p21^CIP1^ and p16^INK4a^ expression. Single nuclei RNA sequencing (snRNA-seq) and proteomics were then performed to define the molecular signature of ovarian cellular senescence in an unbiased manner. With this multi-omics strategy, we identified 26 unique targets that overlap between the tissue transcriptome and secreted proteome. We validated the physiologic relevance of a subset of these proteins by mapping their expression back onto native human ovarian tissue. Overall, our findings reveal a novel and robust senotype of cellular senescence in the human ovary which can enhance our understanding of aging, disease, and the development of therapeutic applications.

**Figure 1.**
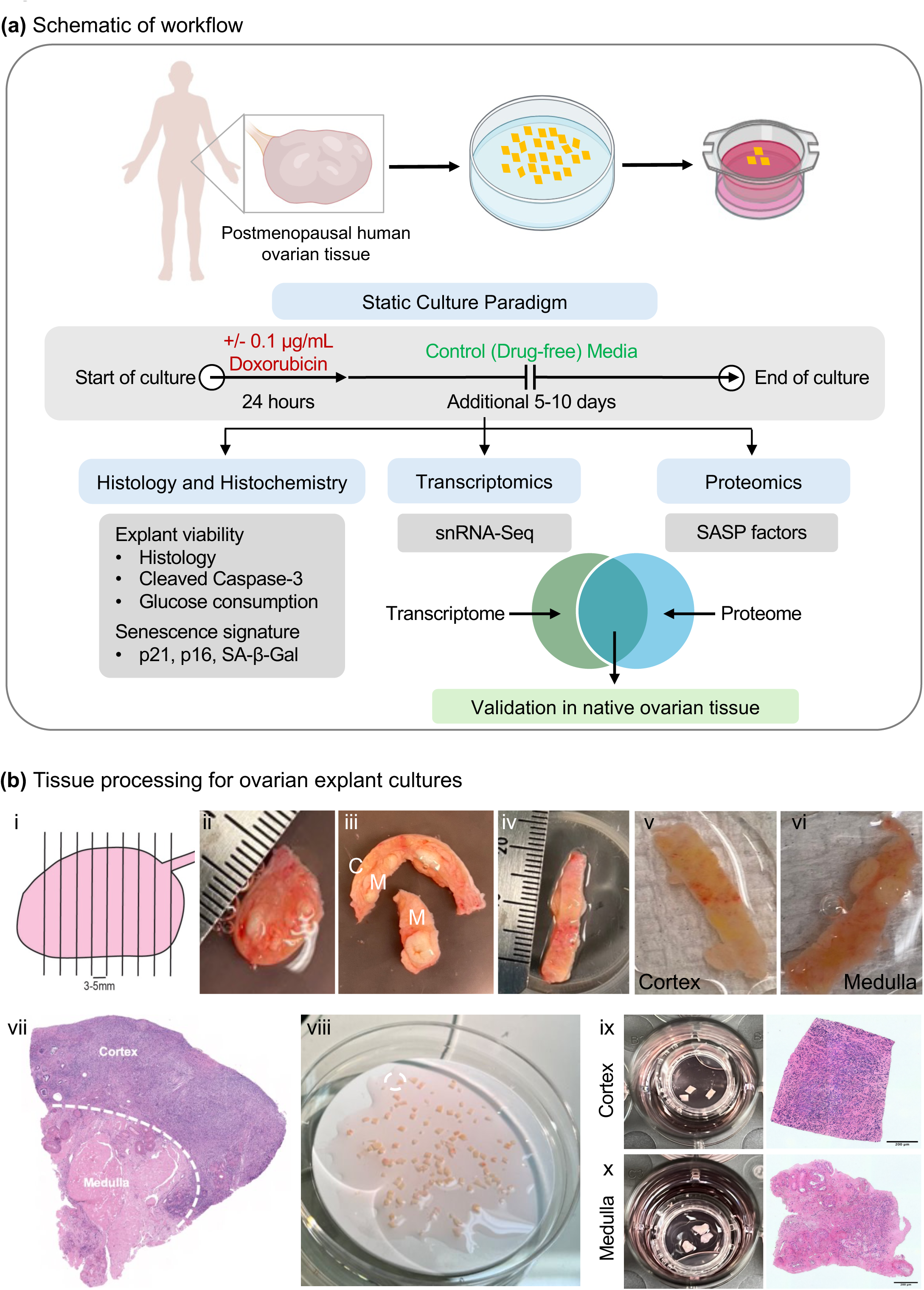
Workflow and tissue processing for the doxorubicin-induced model of cellular senescence in human postmenopausal ovarian explants. (a) Schematic detailing the workflow. Ovarian tissue was obtained from postmenopausal females, processed into explants, and cultured on transwells according to our static culture paradigm. Explants were assessed histologically for viability and senescence markers. Transcriptomics of cultured explants was performed using Single Nuclei Sequencing (snRNA-Seq) and proteomics of conditioned media was performed to analyze SASP factors. A merged signature overlapping between the tissue transcriptome and conditioned media proteome was identified. Key candidates from the merged transcriptomic/proteomic signature were then mapped back onto native postmenopausal ovaries. (b) Tissue processing for ovarian explant cultures (i) Ovaries were sectioned into 3-5 mm thick slices, 1-2 of which are received in lab (ii). (iii) Ovarian sections were then processed into a smaller piece containing both cortex and medulla. (iv) The smaller piece was placed cortex –side –up on a Stadie-Riggs tissue slicer to obtain 500 µm thin slices of the cortex (v) and medulla (vi). (vii) Shows a histological section of an ovarian piece with an outer cortex and inner medulla. (viii) The cortex and medulla slices were processed separately into 1mmx 1mm squares that were cultured as explants on transwells (ix and x; with corresponding histology of a cortex and medulla explant). Scale bars correspond to 200 µm.

## Materials and Methods

### Human ovarian tissue acquisition

De-identified human ovarian tissue was obtained from the Northwestern University Reproductive Tissue Library (NU-RTL) under Institutional Review Board approved protocols (STU00215770, STU00215938). Ovaries were obtained from females aged 50 to 70 years old (Supplemental Table 1; Average 61.3 ± 6.2 years) undergoing bilateral salpingo-oophorectomies and/or total laparoscopic hysterectomies (Supplementary Table 1). Females with BRCA mutations, diagnoses of endometriosis, ovarian neoplasia, complex ovarian cysts, adnexal masses, or a history of breast cancer, radiotherapy, and chemotherapy were excluded. Upon collection, the tissue was divided into cross sections (3-5 mm thick) that were generated perpendicular to the long axis of the ovary (Figure 1b.i) (Devrukhkar et al., 2023). In the absence of significant gross pathology as assessed by a certified gynecologic pathologist, up to two ovarian cross-sections were designated for research (Figure 1b.ii) and transported to the laboratory on ice in ORIGIO^®^ Handling^TM^ IVF medium (Cooper Surgical Inc., Trumbull, CT, USA).

### Ovarian tissue processing and explant culture

Ovarian tissue was kept in the ORIGIO^®^ Handling^TM^ IVF medium for all the processing steps which were performed at room temperature (Anvari et al., 2023). The gross ovarian tissue slice was cut to remove the inner medulla, and a Stadie-Riggs tissue slicer (Thomas Scientific, Chads Fort Township, PA, USA) was then used to generate 500 µm-thick slices of tissue (Figure 1b.iii-vi). As confirmed by histology (Figure 1b.vii), the first slice was cortex-enriched tissue, whereas the last slice was medulla-enriched tissue. The cortex and medulla slices were processed separately and cut into 1mm x 1mm x 0.5 mm pieces with a scalpel (Figure 1b.viii). Ovarian tissue pieces were then transferred into cell culture inserts (Millipore Sigma, Burlington, MA, USA) with three pieces/insert for either the cortex (Figure 1b.ix with corresponding histology) or medulla (Figure 1b.x with corresponding histology). The inserts were then placed in 24-well culture plates with each well containing 400 µL of pre-equilibrated growth media. Growth media was made with MEM Alpha + GlutaMAX (Thermo Fisher Scientific, Waltham, MA, USA) supplemented with 20 mIU/mL recombinant Follicle Stimulating Hormone (Gonal F® RFF Redi-ject®, Rockland, MA, USA), 1 mg/mL fetuin (Sigma-Aldrich, St. Louis, MO, USA), 1 μg/mL Insulin-Transferrin-Sodium Selenite (Thermo Fisher Scientific, Waltham, MA, USA), and 3 mg/mL human serum albumin (Cooper Surgical Inc., Trumbull, CT, USA). This media was prepared with or without doxorubicin hydrochloride (Tocris Bioscience^TM^, Minneapolis, MN, USA) at concentrations of 0, 0.1, and 1 µg/mL depending on the experiment.

Explants were cultured at 37°C in a humidified atmosphere of 5% CO_2_ according to the static culture paradigm (Figure 1a). For doxorubicin treatment, explants were cultured in the compound for 24 hours followed by culture in a doxorubicin-free medium for up to 2 or 5 and 9 additional days for short-term and long-term culture studies, respectively. Explants cultured without doxorubicin throughout the culture served as controls. For short term cultures, complete media changes were performed on days 1 and 3. For long-term cultures, complete media changes were performed on day 1 at the end of doxorubicin treatment followed by half-media changes every other day. Conditioned media was saved throughout culture for downstream analysis. Following the culture period, the explants were either fixed or snap frozen and used for downstream analyses as described. For the SASP analysis, additional steps were taken to ensure minimal background from protein supplements in the conditioned media (Anvari & Duncan, 2023). On day 10, a complete media change was performed with two washes with basal medium without protein supplements (MEM Alpha + GlutaMAX) in the original culture plate. The explants were separated from the inserts by removing the mesh bottom with sterile forceps, and then they were washed in basal medium. Explants were then transferred onto new inserts in plates with fresh basal medium and cultured for an additional 24 hours at which point the conditioned media was snap frozen for proteomic analysis.

### Histochemical and immunohistochemical (IHC) analyses

Explants were fixed in Modified Davidsons fixative (mDF) (Electron Microscopy Sciences, Hatfield, PA, USA) at room temperature for 2 hours and then overnight at 4°C. After overnight fixation, the explants were washed in and transferred to 70% ethanol and stored at 4°C until further processing. The explants were then dehydrated in an automated tissue processor (Leica Biosystems, Buffalo Grove, IL, USA), embedded in paraffin, and sectioned (5 µm thickness) with a microtome (Leica Biosystems, Buffalo Grove, IL, USA). To assess tissue architecture, standard Hematoxylin and Eosin (H&E) staining was performed using a Leica Autostainer XL (Leica Biosystems, Buffalo Grove, IL, USA). Tissue sections were cleared with Xylene (Mercedes Scientific, Lakewood Ranch, FL, USA) in three 5-minute incubations and mounted with Cytoseal XYL (Epredia™ Thermo Fisher Scientific, Waltham, MA, USA).

IHC on ovarian explants was performed with the following antibodies: cleaved caspase-3 (CC3), Ki67, p21^CIP1/WAF1^, p16^INK4A^, Superoxide Dismutase 2 (SOD2), Non-Muscle Myosin IIA (MYH9), Lumican, and Periostin (refer to Supplemental Table 2 for all information including source of antibodies, dilutions, and concentrations) according to a previously established protocol by our laboratory (Machlin et al., 2021). Optimization of all antibodies was performed on native postmenopausal ovarian tissue containing both ovarian cortex and medulla along with positive controls (High Grade Serous Ovarian Carcinoma tissue sections) and non-immune controls (Rabbit and Mouse IgG control antibodies, Vector Laboratories Inc., Burlingame, CA, USA) at the same concentrations as the corresponding primary antibody. In brief, slides were cleared using CitriSolv (Decon Labs, King of Prussia, PA, USA), and rehydrated in decreasing concentrations of ethanol. Antigen retrieval was performed by heat induced epitope retrieval (HIER) using Reveal Decloaker 10X at pH 6 (Biocare Medical, Pacheco, CA, USA) for all antibodies except Periostin (HIER at pH 9.0) and Lumican (no antigen retrieval). Slides were incubated in primary antibodies diluted in 3% Bovine serum albumin (BSA, Sigma-Aldrich, St. Louis, MO, USA) in Tris buffered saline (TBS) at 4°C overnight, followed by incubation in secondary antibody (biotinylated goat anti-rabbit, 1:200, or biotinylated goat anti-mouse, 1:200, Vector Laboratories, Burlingame, CA, USA) for 1 hour at room temperature. Antibody detection was performed using a 3’,3’-diaminobenzidine (DAB) Peroxidase Substrate kit (Vector Laboratories, Burlingame, CA) that resulted in a brown precipitate. As soon as a brown staining was visible in the experimental section, the DAB reaction was stopped by transferring slides to distilled water. The slides were then counterstained with hematoxylin (Mercedes Scientific, Lakewood Ranch, FL, USA), cleared with CitriSolv, and mounted with Cytoseal XYL.

Ovarian explant samples were imaged on a RebelScope Imaging System (Discover ECHO Inc., San Diego, CA, USA) using a 40X objective with 200% optical zoom. Image analysis was performed using FIJI/ImageJ (ImageJ2 Version 2.14.0/1.54f, Madison, WI, USA) (Rueden et al., 2017). Images were processed with background subtraction followed by color deconvolution to split the images into Hematoxylin and DAB-only images. These images were converted to binary images and then the following steps were performed: Watershed-> Threshold (to highlight all nuclei in red)-> Analyze particles (Size in pixels =2500-Infinity). For markers with nuclear localization (CC3, p21^CIP1/WAF1^, and p16^INK4A^) the percentage of positive cells was calculated by dividing the number of DAB positive cells by the total number of nuclei, and the data from three images per explant were averaged. For extracellular matrix (ECM) proteins or those with cytosolic localization (SOD2, MYH9, Lumican, Periostin) the percentage of DAB positive area per total area of each explant was reported.

Native ovarian tissue samples were scanned in brightfield with a 20X Plan Apo objective using the NanoZoomer Digital Pathology whole slide scanning system (HT-9600) (Hamamatsu City, Japan) at the University of Washington Histology and Imaging Core. The Digital Image Analysis (DIA) platform Visiopharm Integrator System (VIS; (Ver. 2023.01.1.13563) (Visiopharm, Hørsholm, Denmark) was used to analyze the IHC. Positive staining was detected by binary thresholding. The percent positive staining was calculated by determining the area of the positive stain label relative to the whole tissue section area.

### Tissue viability assessment

Glucose levels in conditioned media were measured to assess glucose consumption by tissues as a measure of tissue viability (Elson et al., 2015; Prill et al., 2014). Conditioned media was thawed and vortexed, and glucose levels were measured using a GlucCell^®^ glucometer (CESCO Bioengineering, Marina Del Rey, CA, USA) according to manufacturer’s instructions. Baseline measurements on control and doxorubicin-containing media were performed on media samples prior to culture. Glucose levels were then plotted as a mean value from all wells per condition for each participant.

### Senescence Associated Beta-Galactosidase (SA-β-Gal) assay

SA-β-Gal activity was evaluated in frozen tissue sections. Ovarian tissue explants were first embedded in Tissue Tek® optimum cutting temperature (OCT) compound (Sakura® Finetek, VWR, Torrance, CA, USA) and frozen over a mixture of 2-methylbutane (Fisher Scientific, Hampton, NH, USA) and dry ice. The frozen blocks were stored at -80°C until further use. Cryosections of 10 µm thickness were obtained through the Pathology Core Facility (Northwestern University). SA-β-Gal activity was assessed using a Senescence Detection Kit (Biovision, Milpitas, CA, USA) according to the manufacturer’s instructions. In brief, the sections were allowed to equilibrate at room temperature for one minute, fixed with fixative solution for 5 minutes, washed twice in 1X phosphate buffered saline (PBS) (Fisher Scientific, Hampton, NH, USA), and incubated overnight (15 hours) in the SA-β-Gal staining solution at 37°C. The slides were then washed in PBS, counterstained with Nuclear Fast Red (Vector Laboratories, Newark, CA, USA) for 5 minutes and mounted in aqueous mounting medium (CC/mount^TM^, Sigma-Aldrich, St. Louis, MO, USA). The stained tissues were imaged under the RebelScope Imaging System using a 40X objective with 200% optical zoom. The presence of blue staining was considered indicative of SA-β-Gal positivity.

### Single nuclei RNA sequencing (snRNA-seq)

Human ovary explants were transported on dry ice from Northwestern University to the Buck Institute for Research on Aging and transferred to -80°C upon arrival. Explants were transferred into a pre-chilled 1.5 mL Eppendorf tube and immediately dissociated into single nuclei suspension using the 10X Genomics nuclei isolation kit (PN: 1000494) using a 15-minute lysis with two resuspension washes. A cordless motor pestle (VWR, Radnor, PA, USA) was used to dissociate the tissue into a single nuclei suspension. Final nuclei concentration was determined using the countess II automated cell counter (Thermo Fisher, Waltham, MA, USA) with propidium iodide (Invitrogen, Waltham, MA, USA). Single-cell libraries were then prepared using the Chromium Next Gem Automated Single 5’ Library and Gel Bead Kit v.2 (PN100290) on a Chromium Connect robot (PN1000171), following the manufacturer’s instructions. The cDNA and final gene expression libraries were quantified using a tape station (Agilent Technologies Inc., Santa Clara, CA, USA) and submitted for sequencing.

### Processing and analysis of snRNA-seq data

Data processing was performed using 10x Genomics Cell Ranger v6.1.2 pipeline. The “cellranger count” was used to perform transcriptome alignment, filtering, and UMI counting from the FASTQ (raw data) files. Alignment was done against the human genome GRCh38-2020-A. Cell numbers after processing were: 10-day explants: doxorubicin-treated cortex 7,207 cells, control cortex 7,980 cells, doxorubicin-treated medulla 3,774 cells, and control medulla 5,324 cells; 6-day explants: doxorubicin-treated cortex 24,702 cells, control cortex 11,332 cells, doxorubicin-treated medulla 22,852 cells, and control medulla 12,808 cells.

Downstream analyses were performed in R (version 4.2.0), primarily using the Seurat R package (version 4.1.1)(Hao et al., 2021; Satija et al., 2015) and custom analysis scripts. First, we executed a quality control step that removed the cells containing >10% mitochondrial RNA and <=250 genes/features. The doublet cells were identified and removed from the downstream analysis by using the DoubletFinder R package (version 2.0.3)(McGinnis et al., 2019) with parameters PCs=1:30, pN=0.25, and nExp=7.5%. A total of 22,055 cells from 10-day explant cultures and 71,694 cells from 6-day explant cultures remained for subsequent analysis. Raw RNA counts were first normalized and stabilized with the SCTransform v2 function (SCT), then followed by the RPCA integration workflow for joint analysis of single-cell datasets. In doing so, the top 3,000 highly variable genes/features among the datasets were used to run SCT; and then 3,000 highly variable genes/features and the 30 top principal components (PCs) with k.anchor=5 were used to find “anchors” for integration. The clustering step was executed by using the 30 top PCs summarizing the RNA expression of each cell with a resolution parameter of 0.8.

To identify putative cell types, singleR R package with (version 2.0.0)(Aran et al., 2019) was used with the reference dataset of human primary cell atlas data (HPCA). Cell type annotation results from singleR were further refined by checking the manually curated marker gene list for main cell types present in the human ovary (Supplemental Data 3) (Fan et al., 2019; Ernst Lengyel et al., 2022; Wagner et al., 2020). The correlation and enrichment analyses of marker gene expression were conducted to assist in determining the cell types. Differential expression gene (DEG) analyses were done by functions in Seurat PrepSCTFindMarkers then FindAllMarkers/FindMarkers functions with MAST algorithm (Finak et al., 2015) . For the overall analyses and each cell type, the comparisons of DEGs of doxorubicin against control in either cortex or medulla ovarian tissue (Supplemental Data 4 and 5) were calculated by using the FindMarker function with parameter min.pct=0.1 and logFC=0.2. Rank-Rank Hypergeometric Overlap (RRHO) analysis (Cahill et al., 2018; Plaisier et al., 2010) was performed by using the RRHO2 R package (version 1.0) to compare the differential expression patterns between doxorubicin and control of cortex vs medulla tissues. The ranks of the genes in the two gene lists were determined by calculating -log10(adj.pvalue)*Log2FC.

Following differential expression, Ingenuity Pathway Analysis (IPA, Qiagen) was used to discover changes in enriched pathways in each comparison. DEGs with adjusted p-values < 0.05 and |Log2FC|> 0.2 were incorporated into the IPA canonical pathway analysis.

The degree of cellular senescence was quantified utilizing the AUCell R package (Aibar et al., 2017) which scores cell activity based on gene expression profiles. We applied multiple gene sets associated with cellular senescence, sourced from various peer-reviewed databases and publications, as well as SASP factors from this study (“Buck ovary SASP Cortex and Medulla”) (Supplemental Data 6) (Ansere et al., 2021; Nathan Basisty et al., 2020; de Magalhaes et al., 2023; de Magalhães et al., 2009; Gao et al., 2023; Kiss et al., 2020; Landry et al., 2022; E. Lengyel et al., 2022; Milacic et al., 2023; Saul et al., 2022; Shen et al., 2019; Thomas et al., 2022), to ensure comprehensive coverage and robustness of the senescence scoring. The two-sided Student’s t-test was used to compare the difference in senescence scores between doxorubicin and control groups. Statistical significance was established at p-value < 0.05.

### Proteomic sample preparation and analysis

To assess SASP factors in conditioned media by proteomics, 400 μL of conditioned media from human ovarian cortex and medulla explant tissue cultures in both doxorubicin treatment and control conditions was concentrated to ∼30 µL with 0.5 mL 3 kDa filters (Millipore Sigma, Burlington, MA, USA). Aliquots of concentrated secretome (15 µL) for each sample were reduced using 20 mM dithiothreitol in 50 mM triethylammonium bicarbonate buffer (TEAB) at 50°C for 10 min, cooled to room temperature (RT) and held at RT for 10 min, and alkylated using 40 mM iodoacetamide in 50 mM TEAB at RT in the dark for 30 min. Samples were acidified with 12% phosphoric acid to obtain a final concentration of 1.2% phosphoric acid. S-Trap buffer consisting of 90% methanol in 100 mM TEAB at pH ∼7.1, was added and samples were loaded onto the S-Trap micro spin columns. The entire sample volume was spun through the S-Trap micro spin columns at 4,000 x *g* and RT, binding the proteins to the micro spin columns. Subsequently, S-Trap micro spin columns were washed twice with S-Trap buffer at 4,000 x *g* at RT and placed into clean elution tubes. Samples were incubated for one hour at 47°C with 2 µg of sequencing grade trypsin (Promega, San Luis Obispo, CA) dissolved in 50 mM TEAB. Afterwards, trypsin solution was added again at the same amount, and proteins were digested overnight at 37°C.

Proteolytic peptides were sequentially eluted from micro S-Trap spin columns with 50 mM TEAB, 0.5% formic acid (FA) in water, and 50% acetonitrile (ACN) in 0.5% FA. After centrifugal evaporation, samples were resuspended in 0.2% FA in water and desalted with Oasis 10 mg Sorbent Cartridges (Waters, Milford, MA). The desalted protein lysates were then subjected to centrifugal evaporation and re-suspended in 30 µL of 0.2% FA in water. Finally, indexed Retention Time standard peptides (iRT, Biognosys, Schlieren, Switzerland) (Escher et al., 2012) were spiked into the samples according to manufacturer’s instructions.

### Mass spectrometric analysis

LC-MS/MS analyses were performed on a Dionex UltiMate 3000 system online connected to an Orbitrap Eclipse Tribrid mass spectrometer (both Thermo Fisher Scientific, San Jose, CA). The solvent system consisted of 2% ACN, 0.1% FA in water (solvent A) and 98% ACN, 0.1% FA in water (solvent B). Proteolytic peptides (2 µL of 1:40-diluted sample) were loaded onto an Acclaim PepMap 100 C_18_ trap column (75 µm x 20 mm, 3 µm particle size; Thermo Fisher Scientific) for 10 min at 5 µL/min with 100% solvent A. Peptides were eluted on an Acclaim PepMap 100 C_18_ analytical column (75 µm x 50 cm, 3 µm particle size; Thermo Fisher Scientific) at 300 nL/min using the following gradient of solvent B: 2% for 10 min, linear from 2% to 20% in 125 min, linear from 20% to 32% in 40 min, up to 80% in 1 min, 80% for 9 min, and down to 2% in 1 min. The column was equilibrated with 2% of solvent B for 29 min, with a total gradient length of 215 min.

All samples were acquired in data-independent acquisition (DIA) mode (Bruderer et al., 2017; Collins et al., 2017; Gillet et al., 2012). Full MS spectra were collected at a resolution of 120,000 (AGC target: 3e6 ions, maximum injection time: 60 ms, 350-1,650 m/z), and MS2 spectra at a resolution of 30,000 (AGC target: 3e6 ions, maximum injection time: Auto, NCE: 27, fixed first mass 200 m/z). The isolation scheme consisted in 26 variable windows covering the 350-1,650 m/z range with an overlap of 1 m/z (Supplemental Data 7) (Bruderer et al., 2017).

### DIA data processing and analysis

DIA data was processed in Spectronaut (version 17.6.230428.55965) using directDIA. Data was searched against a human database containing all UniProt-SwissProt entries extracted on 06/30/2023 (20,423 entries). Trypsin/P was set as the digestion enzyme and two missed cleavages were allowed. Cysteine carbamidomethylation was set as a fixed modification while methionine oxidation and protein N-terminus acetylation were set as dynamic modifications. Data extraction parameters were set as dynamic and non-linear iRT calibration with precision iRT was selected. Identification was performed using 1% precursor and protein q-value. iRT profiling was selected. Quantification was based on the peak areas of extracted ion chromatograms (XICs) of the 3-6 best fragment ions per precursor ion, and q-value sparse data filtering was applied. Interference correction was selected, and no normalization was applied. Differential protein abundance analysis was performed using an unpaired t-test, and p-values were corrected for multiple testing, using the Storey method (Burger, 2018; Storey, 2002). Protein groups were required with at least two unique peptides. Protein groups with q-value < 0.05 and absolute Log_2_(fold-change) > 0.58 were considered significantly altered (Supplemental Data 8).

Partial least square-discriminant analysis (PLS-DA) of the proteomics data was performed using the package mixOmics n R (version 4.0.2) (Rohart et al., 2017).

### Statistical analysis

All graphs were generated using GraphPad Prism Software Version 9.4.1. Values were represented as mean ± SD. For immunohistochemistry analysis, normality was assessed by the Shapiro-Wilk test and statistical significance was determined using unpaired t-tests and one-way ANOVA, with p values <0.05 considered statistically significant. For transcriptomics data, the two-sided Student’s t-test was used to compare the difference in senescence scores between doxorubicin and control groups and statistical significance was established at p-value < 0.05. For proteomics data, differential protein abundance analysis was performed using an unpaired t-test, and p-values were corrected for multiple testing, using the Storey method (Burger, 2018; Storey, 2002). Protein groups with q-value < 0.05 and absolute Log_2_(fold-change) > 0.58 were considered significantly altered.

## Results

### Development of a human ovarian explant culture model of induced senescence

Post-menopausal human ovarian tissue contains numerous cell types, including fibroblasts, endothelial, epithelial, smooth muscle, and immune cells (E. Lengyel et al., 2022). Therefore, we developed an ovarian tissue explant culture model to interrogate doxorubicin-induced senescence in an intact tissue that maintained this cellular heterogeneity. The post-menopausal ovary exhibits distinct architectural regions consisting of a cellular dense outer cortex and a vascular rich inner medulla (Figure 1b.vii). Therefore, we analyzed cortical and medullary tissues separately. We first determined an optimal dose that induced senescence but did not impact explant viability in culture. Ki67 and cleaved caspase-3 (CC3) are biomarkers of cell proliferation and apoptosis, respectively, which are commonly used to evaluate cell/tissue viability, and the baseline expression of both are low in the post-menopausal ovary (Figures S1a-b). Since apoptosis is one of the main pathways involved in doxorubicin-induced cell death in the ovary (Spears et al., 2019), we used it as a primary marker of viability in our explant model. First, we evaluated tissue morphology by H&E staining and apoptosis levels quantifying CC3 expression in explants exposed to different concentrations of doxorubicin (0, 0.1, and 1 µg/mL) for 24-hours followed by 2 days of culture in doxorubicin-free control medium (Figure S2). Doxorubicin treatment did not impact the tissue morphology of ovarian cortical (Figure S2a) or medullary explants (Figure S2b) nor show any histological evidence of tissue necrosis (nuclear condensation, eosinophilic cytoplasm, pyknotic nuclei) when compared to control tissues (Laronda et al., 2014; Otala et al., 2002). The explant cultures also did not exhibit appreciable cell death on day 1 or day 3 of the culture relative to controls, suggesting our treatment paradigm did not affect explant viability (Figure S2c-S2f). Ultimately, we selected a concentration of 0.1 µg/mL doxorubicin for subsequent long-term cultures based given that this dose has been used in other cell types to induce cellular senescence and the viability we established in our explant culture model (Alimirah et al., 2020; Demaria et al., 2017; Kitada et al., 2019).

For long-term cultures, explants were cultured with or without doxorubicin (0 and 0.1 µg/mL) for 24 hours, followed by an additional 5 and 9-days of culture in a doxorubicin-free medium (Figure 1a). Doxorubicin did not affect gross tissue morphology or show histological evidence of overt tissue necrosis in both cortical or medullary explants as compared to Day 0 and control tissues at both 6 (Figure 2a) and 10 days (Figure 2b) of culture. Additionally, the explants displayed ovarian surface remodeling, a characteristic of wound healing and healthy stroma as seen by the smooth epithelial edge at the later stages of culture as compared to the rough edges at day 0 (Figure 2a & 2b) (Jackson et al., 2009; Laronda et al., 2014). The level of apoptosis in cultured explants was also not significantly changed for both 6- and 10-day cultures, suggesting that the cultured explants were viable (Figures 2c & 2e). We further confirmed tissue viability by assessing metabolic activity via measurement of glucose levels in conditioned media which is commonly used as a non-invasive method to assess viability in long-term cultures (Elson et al., 2015; Prill et al., 2014). Glucose levels in our cultures decreased throughout culture, indicating that the tissues were consuming glucose (Figures 2d & 2f). Taken together, we confirmed ovarian explant viability through a combination of histological, immunohistochemical, and metabolic assays.

**Figure 2.**
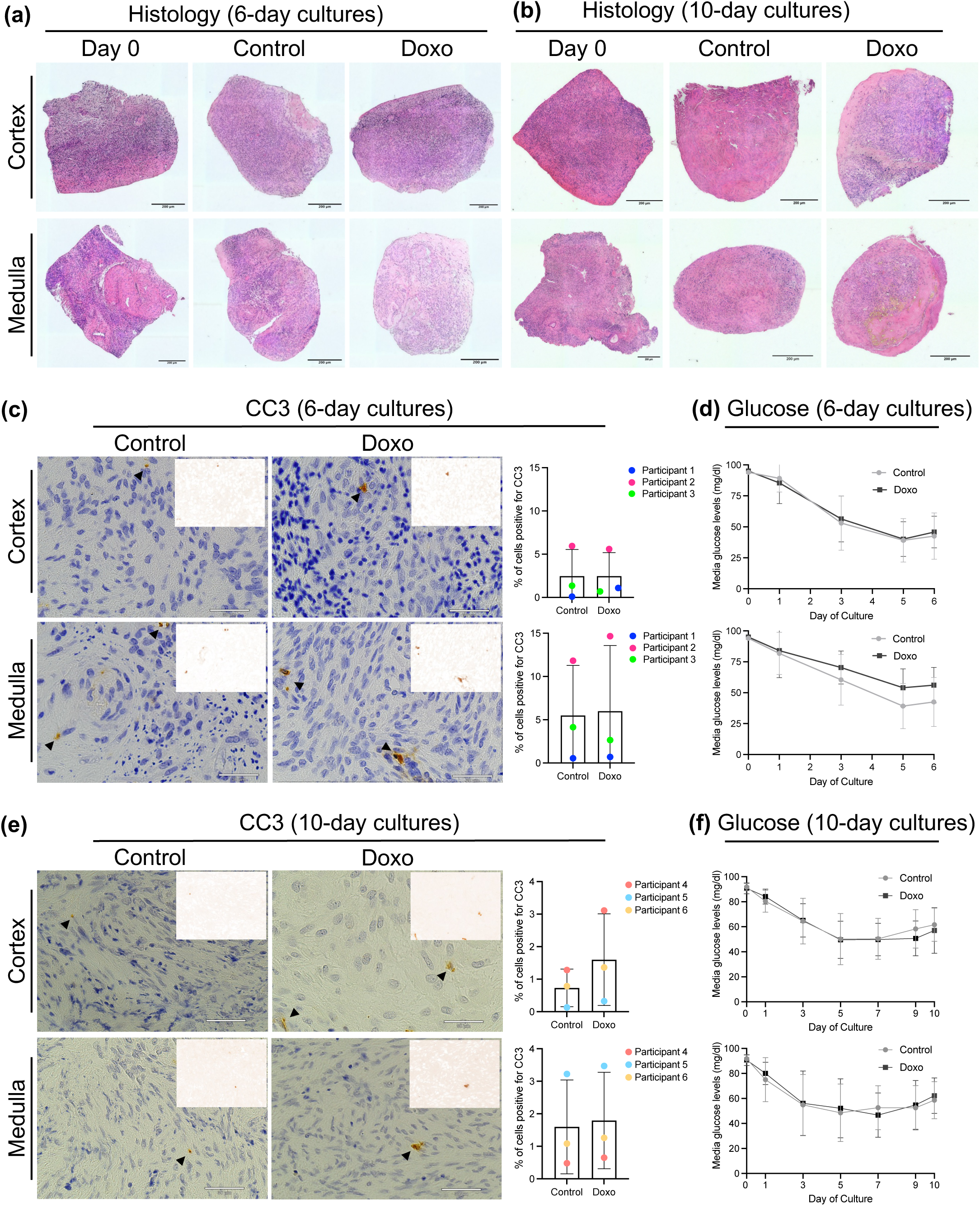
Assessment of ovarian tissue explant viability after 6 and 10 days of culture. (a- b) H&E-stained sections of human ovarian cortex and medulla explants cultured for 6 days (a) and 10 days (b) with 24 hours doxorubicin exposure (0 or 0.1 µg/mL). Histology at Day 0, uncultured tissues is shown for comparison. Explants show no change in gross morphology, no signs of tissue necrosis, and show smoothened edges at 6- and 10 days indicative of wound healing and a healthy stroma (c) Immunohistochemistry for CC3 in the ovarian cortex and medulla explants on day 6 of culture showed low levels of cellular apoptosis (p-value >0.05). (d) Glucose levels measured in conditioned media of the cultured cortex and medulla explants on days 0, 1, 3, 5, and 6 showed decreasing glucose levels in conditioned media, indicating glucose consumption by explants throughout 6 days of culture. Values are represented as a mean of 9 replicate wells per condition (N=3 participants 61, 64, and 65 years old). (e) Immunohistochemistry for CC3 in the ovarian cortex and medulla explants on day 10 of culture showed low levels of cellular apoptosis (p-value >0.05). (f) Glucose levels measured in conditioned media of the cultured cortex and medulla explants on days 0, 1, 3, 5, 7, 9, and 10 show decreasing glucose levels in conditioned media, indicating glucose consumption by explants throughout 10 days of culture. Values are represented as a mean of 9 replicate wells per condition (N=3 participants 50, 53, 62 years old). Insets: Color deconvoluted images highlighted DAB-positive cells in brown color. Statistical significance was determined using an unpaired t-test and and p values <0.05 were considered statistically significant.

### Canonical senescence markers tended to increase with doxorubicin treatment

Although there is no single hallmark of senescent cells, there are several markers that tend to be conserved in different cell types and tissues, including increased senescence-associated beta-galactosidase (SA-β-Gal) activity and p21^CIP1^ and p16^INKa^ expression (Dimri et al., 1995; Kumari & Jat, 2021; Suryadevara et al., 2024). Of note, these markers increase in response to doxorubicin in several cell and tissue types (Alimirah et al., 2020; Altieri et al., 2016; Bielak-Zmijewska et al., 2014; Hou et al., 2019; Piegari et al., 2013). We observed more pronounced SA-β-Gal signal in doxorubicin-treated tissues compared to the controls (Figure S3a). With respect to p21^CIP1^ and p16^INKa^, we noted minimal p21^CIP1^ expression in native postmenopausal ovarian tissue (Figures S1b & S1d), whereas p16^INK4a^ was expressed in clusters throughout the ovarian cortex and medulla (Figures S1c & S1d). In our induced senescence model, p21^CIP1^ and p16^INK4a^ expression tended to increase with doxorubicin in both 6-day and 10-day cultured explants, but there was marked heterogeneity among individual participants (Figures S3b-e).

### Single nuclei RNA sequencing highlights distinct senescence-induced transcript signatures in the ovarian cortex and medulla

To gain deeper and unbiased molecular insight into an ovarian senescence signature induced by doxorubicin treatment, we performed single nuclei RNA sequencing (snRNA-seq) to assess the transcriptomic signature of cultured ovarian explants (Figure 1a). A total of 71,694 cells was profiled from 6-day explants (cortex control=11,332, cortex doxorubicin treated=24,702, medulla control=12,808, medulla doxorubicin treated=22,852), and a total of 22,055 cells were profiled from 10-day explants (cortex control=7,980, cortex doxorubicin treated=7,207, medulla control=5,324, medulla doxorubicin treated=3,744) as shown in the UMAP (Figure 3a). We utilized eleven senescence gene sets compiled from peer-reviewed databases and publications to calculate a cellular senescence score based on gene expression fold change profiles across 6- and 10-day cortex and medulla explants (Figure 3b). Based on this analysis, the 10-day explants had consistently higher cellular senescence scores relative to the 6-day explants in both the cortex and medulla (Figure 3b). Interestingly the cortex had a much higher senescence score than the medulla which was driven by distinct gene sets (cortex: 4, 11, 2, 1, 3, 6; medulla: 9, 5, 7, 10, 8) suggesting an ovarian compartment-specific difference in response to senescence induction (Figure 3c; Figure S4a-b).

**Figure 3.**
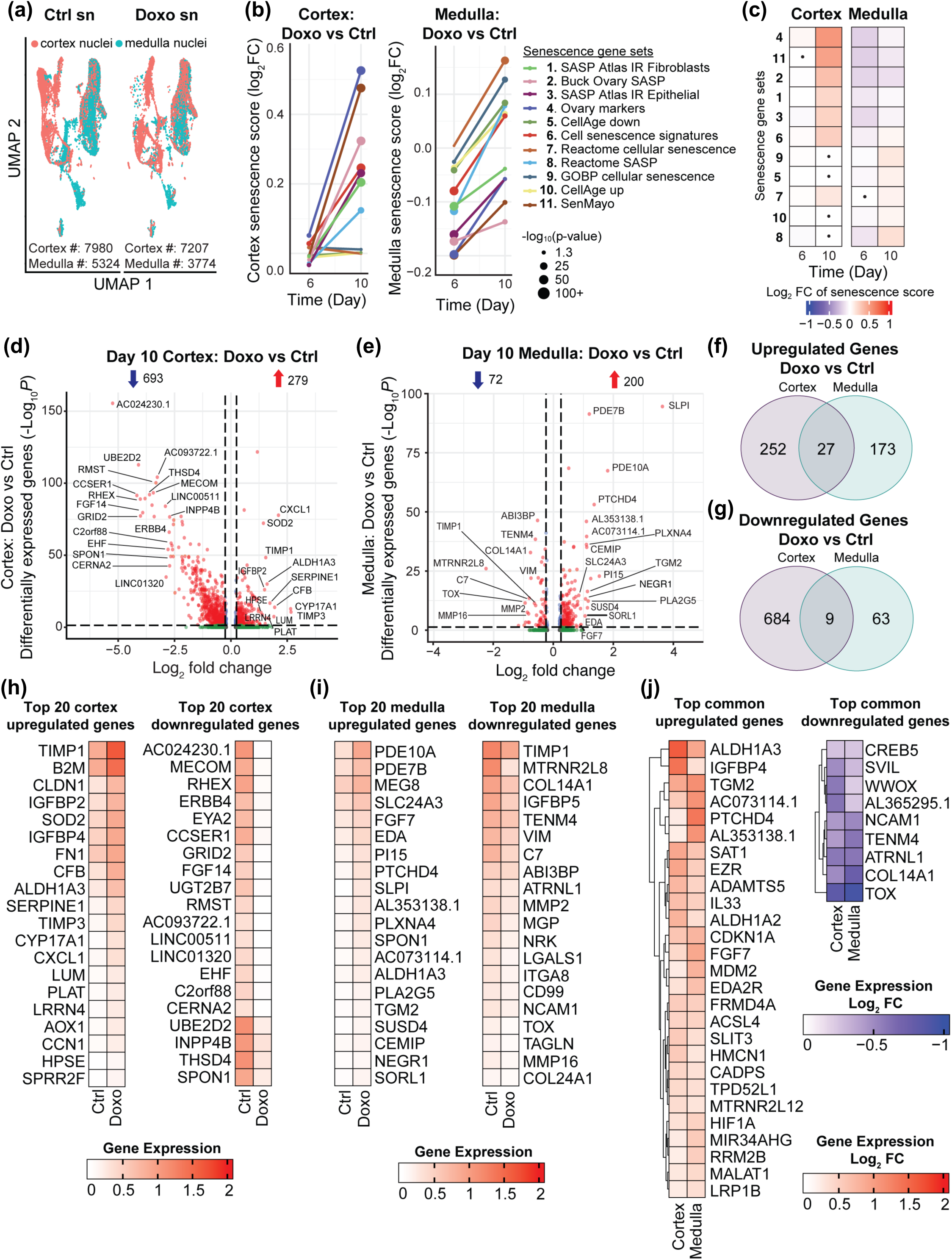
snRNA-seq highlights differences in senescence induction in ovarian cortex and medulla. (a) UMAP plot of single-nuclei (sn) transcriptomes of explant ovarian tissue divided into cortex (pink) and medulla (green) after 10-day treatment with doxorubicin (Doxo) or without doxorubicin (Ctrl). (b) Trajectory of cellular senescence scores across day 6 and 10 in ovarian cortex and medulla tissue. X-axis is the days post doxo treatment. Y-axis is the Log2FC of senescence scores in the doxo-treated sample relative to the untreated control. The size of the dot is the –log10 transformed p-value from two-sided t-test. (c) Heatmaps depicting senescence scores across the cortex and medulla over 6- and 10-day doxo treatment. These scores were calculated using 11 gene sets associated with cellular senescence. The heatmap values represent log2 fold change of senescence scores in doxo-treated cells relative to untreated controls. Non-significant differences are indicated with a dot. Statistical significance was assessed using two-sided t-test, with a threshold of p<0.05. (d) A volcano plot of differentially expressed genes (DEGs) in day 10 cortex (Doxo vs Ctrl). DEGs (including Log2FC and p-value) were calculated by the MAST method. The Benjamini-Hochberg method was used for multiple comparison adjustments. p-value (adj. p) cutoff is <0.05, and Log2FC cutoff is >0.25. (e) A volcano plot of differentially expressed genes (DEGs) in day 10 medulla (Doxo vs Ctrl). DEGs (including Log2FC and p-value) were calculated by the MAST method. The Benjamini-Hochberg method was used for multiple comparison adjustments. p-value (adj. p) cutoff is <0.05, and Log2FC cutoff is >0.25. (f) Venn diagram showing the overlap of upregulated DEGs between day 10 cortex and medulla (Doxo vs ctrl). (g) Venn diagram showing the overlap of downregulated DEGs between day 10 cortex and medulla (Doxo vs ctrl). (h-i) Heatmaps depicting the top 20 absolute gene expression of up- and downregulated DEGs in cortex and medulla comparing Ctrl and Doxo-treated expression. (j) Heatmaps depicting the Log2FC of the 27 shared upregulated DEGs, and the 9 downregulated DEGs in the cortex and medulla comparing Ctrl and Doxo-treated expression.

Given the time-dependent increase in cellular senescence response, further analyses were restricted to the 10-day explants. At a global level doxorubicin treatment relative to the control resulted in 693 downregulated and 279 upregulated differential expressed genes (DEGs) in the cortex, and 72 downregulated and 200 upregulated DEGs in the medulla (Figures 3d-e). Of the upregulated DEGs, 27 were shared between the cortex and medulla (10% of cortex DEGs and 14% of medulla DEGs) (Figure 3f, Supplemental Data 4 & 5). Of the downregulated DEGs, 9 were shared between the cortex and medulla (1% of cortex DEGs and 13% of medulla DEGs) (Figure 3g, Supplemental Data 4 & 5). We identified the top 20 upregulated and downregulated DEGs in the cortex (Figure 3h) and medulla (Figure 3i), showing a distinct senescence regional signature. However, we also identified the key upregulated and downregulated DEGs common between the cortex and medulla (Figure 3j). Importantly, cyclin-dependent kinase inhibitor 1A (CDKN1A), better known as p21, a common marker of senescence was equally upregulated in both regions (Figure 3j), consistent with the trend in IHC staining observed in Figures S3b & S3d. Based on Ingenuity Pathway Analysis (IPA), doxorubicin altered genes in the cortex that are enriched in pathways associated with inflammation, fibrosis, oxidative stress, and immune responses, in particular IL-6 and HMGB1 signaling, which are common hallmarks of senescence (Figure S5a). However, there was a reversal in activated pathways between the two regions such as the unfolded protein response and the immunogenic cell death signaling pathways (Figure S5a & b). These were among the highest activated pathways in the cortex but the most inactivated pathway in the medulla (Figure S5a-b). Collectively these results implicate a compartment-specific response to senescence induction in the human ovary.

### snRNA-seq reveals cell type-specific senescence signatures after doxorubicin treatment

A major strength of our model is the ability to induce cellular senescence within the context of an intact tissue with cellular heterogeneity. To further understand the senescence response at a cellular level, we analyzed cellular composition in day 10 cultured explants using the Uniform Manifold Approximation and Projection (UMAP) algorithm to plot the 8 identified transcriptionally distinct cell clusters based on cell type-specific markers (Figure 4a, Supplemental Data 3). Processing the cortex and medulla separately in our paradigm allowed us to observe proportional cell type differences between these compartments. Except for epithelial cells in the cortex, the proportion of each cell type was similar in each region and experimental condition (Figure 4b). As expected, given the presence of the ovarian surface epithelium, epithelial cells were greatly enriched in the control cortex relative to the medulla. This cell type, however, was highly sensitive to doxorubicin treatment and significantly decreased following treatment (Figure 4b). A senescence score was calculated to determine the enrichment of senescence genes across the eight distinct cell clusters in the cortex and medulla, permitting us to localize potential cell types driving the senescence signature we identified in the cortex and medulla (Figure 3). Our results revealed that epithelial and stromal cells (Stroma 1) in the cortex (Figure 4c & S4c-d) and stromal cells (Stroma 1) in the medulla (Figure 4d & S4e) exhibited the highest senescence scores when compared to the overall tissue score.

**Figure 4.**
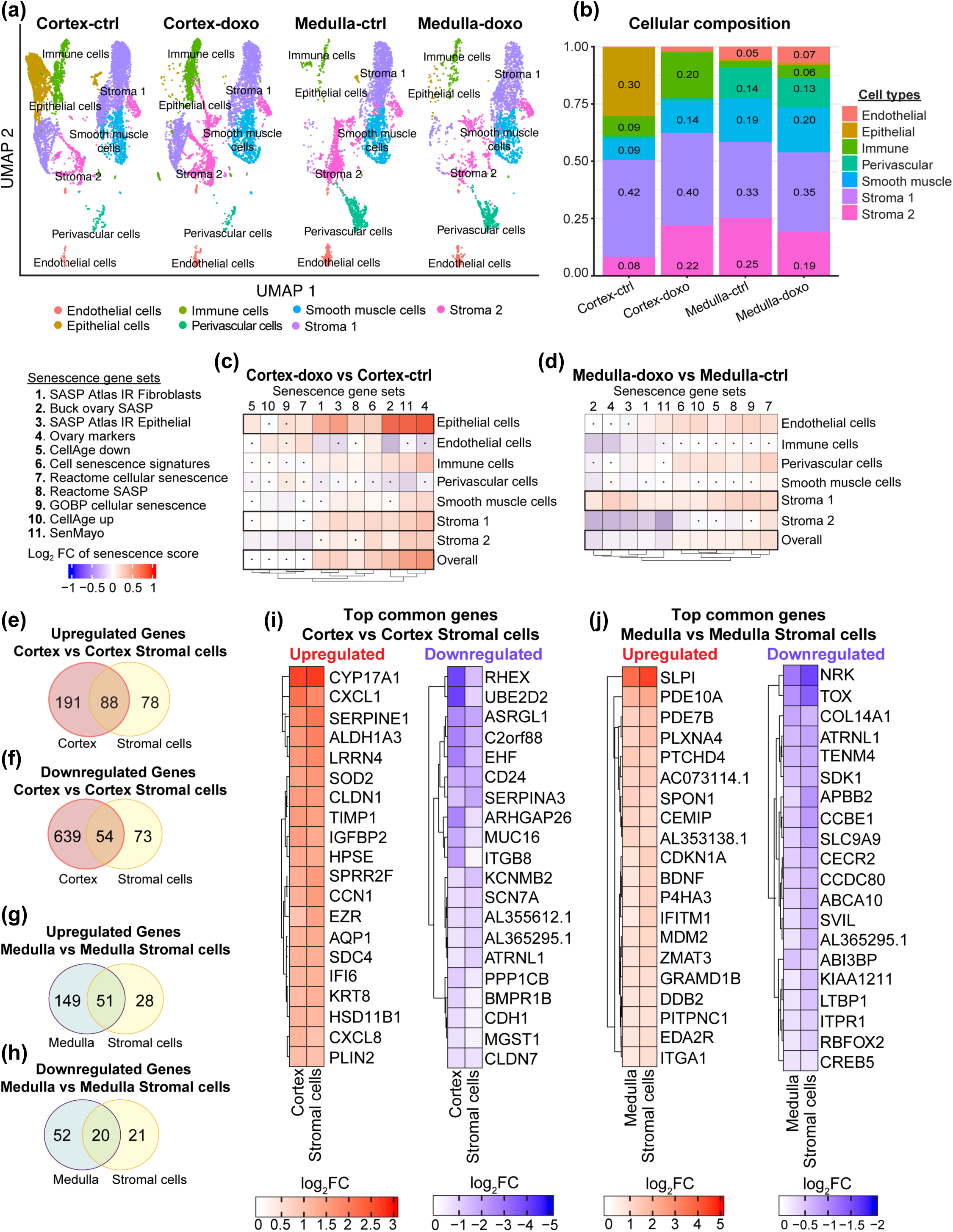
Cell composition analysis by snRNA-seq reveals cell types driving senescent signature in the ovary. (a) A UMAP plot of 10-day single-nuclei transcriptomes of explant ovarian tissue divided into cortex and medulla after doxorubicin (Doxo) treated and untreated (Ctrl). Cells were resolved into 8 distinct cell types. (b) A stacked bar chart showing the quantification of the relative abundance of each cell type in cortex and medulla treated with (Doxo) and without doxorubicin (Ctrl) expressed by percent. (c-d) Heatmaps depicting senescence scores across different cell types within the ovarian cortex (c) and medulla (d) after 10-day treatment with doxorubicin. These scores were calculated using 11 gene sets associated with cellular senescence. The heatmap values represent log2 fold changes of senescence scores in doxo-treated cells relative to untreated controls. Non-significant differences are indicated with a dot. Statistical significance was assessed using a two-sided t-test, with a threshold of p<0.05 for comparing doxo-treated cells to controls within the same cell type. (e) Venn diagram showing the overlap of upregulated DEGs between day 10 cortex and cortex stromal cells (Stroma 1) (f) Venn diagram showing the overlap of downregulated DEGs between day 10 cortex and cortex stromal cells (Stroma 1). (g) Venn diagram showing the overlap of upregulated DEGs between day 10 medulla and medulla stromal cells (Stroma 1). (h) Venn diagram showing the overlap of downregulated DEGs between day 10 medulla and medulla stromal cells (Stromal 1). (i-j) Heatmaps depicting the top 20 Log2FC up- and downregulated shared DEGs between cortex and cortex stromal cells (i) and medulla and medulla stromal cells (j).

Although cortical epithelial cells had the highest senescence score in the cortex, only 15 upregulated genes (Figure S6a) and 10 downregulated genes (Figure S6b) were common with the overall cortex DEGs. Nonetheless, 3 DEGs out of the top 20 upregulated cortex genes are shared with the epithelial cells, PLAT, CFB, and SOD2 (Figure 3h & S6c). Whereas 5 DEGs out of the top 20 downregulated cortex genes are shared with the epithelial cells, A2093722.1, A2024230.1, THSD4, EYA2, and CCSER1 (Figure 3h & S6d). However, cortex stromal cells (Stroma 1) contributed more shared DEGs with the overall cortex DEGs. Of the upregulated genes, 88 were shared (32% of overall cortex DEGs and 53% of stromal DEGs) (Figure 4e), whereas 12/20 of the top 20 upregulated cortex DEGs were shared including TIMP1, SOD2, and SERPINE 1 (Figure 3h & 4i). There were 54 shared downregulated genes between cortex and cortex stroma (8% of overall cortex DEGs and 43% of stromal DEGs) (Figure 4f & 4i). In the medulla, there was a greater overlap of DEGs with medulla stromal cells (Stroma 1). There were 51 shared upregulated genes (26% of overall medulla DEGs and 65% of stromal DEGs) (Figure 4g), whereby 9/20 of the top 20 upregulated medulla DEGs were shared including SLPI, PDE10A, and SPON1 (Figure 3i & 4j). Between the medulla and medulla stroma, 20 downregulated genes were shared (28% of overall medulla DEGs and 49% of stromal DEGs) (Figure 4h & 4j). These results suggest that epithelial and stromal cells (Stroma 1) in the cortex and stromal cells (Stroma 1) in the medulla might be driving the senescence signature in these regions.

Ingenuity Pathway Analysis (IPA) provided an overview of enriched pathways in the stromal cells of the cortex and medulla revealing contrasting activated pathways (Figures S7a & S7b). The top significantly activated pathway in the cortex stroma was ‘Integrin signaling’, which regulates cell-cell and cell-extracellular matrix adhesion, including cellular proliferation/migration and activation/release of cytokines (Figure S7a). The top significantly activated pathway in the medulla stroma was ‘EIF2 signaling’, which regulates mRNA translation and controls proteome expression of downstream pathways such as the Integrated Stress Response (ISR) and inflammatory production of cytokines (Figure S7b). These results unveil the cell type specific heterogeneity in response to doxorubicin-induced senescence in human ovaries.

### Proteomic profiling of human ovarian explants identified SASP factors upon senescence induction by doxorubicin

Senescent cells are characterized by production and release of a prominent secretome known as the Senescence Associated Secretory Phenotype (SASP) (Di Micco et al., 2021; Hernandez-Segura et al., 2018; Herranz & Gil, 2018; Kumari & Jat, 2021; Wiley & Campisi, 2021). We were able to profile the SASP in our model by assessing the conditioned media using proteomics (Figure 5a, Supplemental Table 1). On day 10 of culture, tissues were thoroughly washed with serum-free basal media (alpha-MEM) and transferred to inserts in a clean plate with pre-equilibrated serum-free basal media (Figure 5a). The conditioned media was then collected after 24 hours for proteomic analysis (6 replicate media wells/condition, except “medulla control” that had media from 4 replicate wells (400 µL of conditioned media/well). Proteins from the conditioned media were concentrated and prepared for proteomic analysis as described using directDIA to identify SASP factors (Figure 5a).

**Figure 5.**
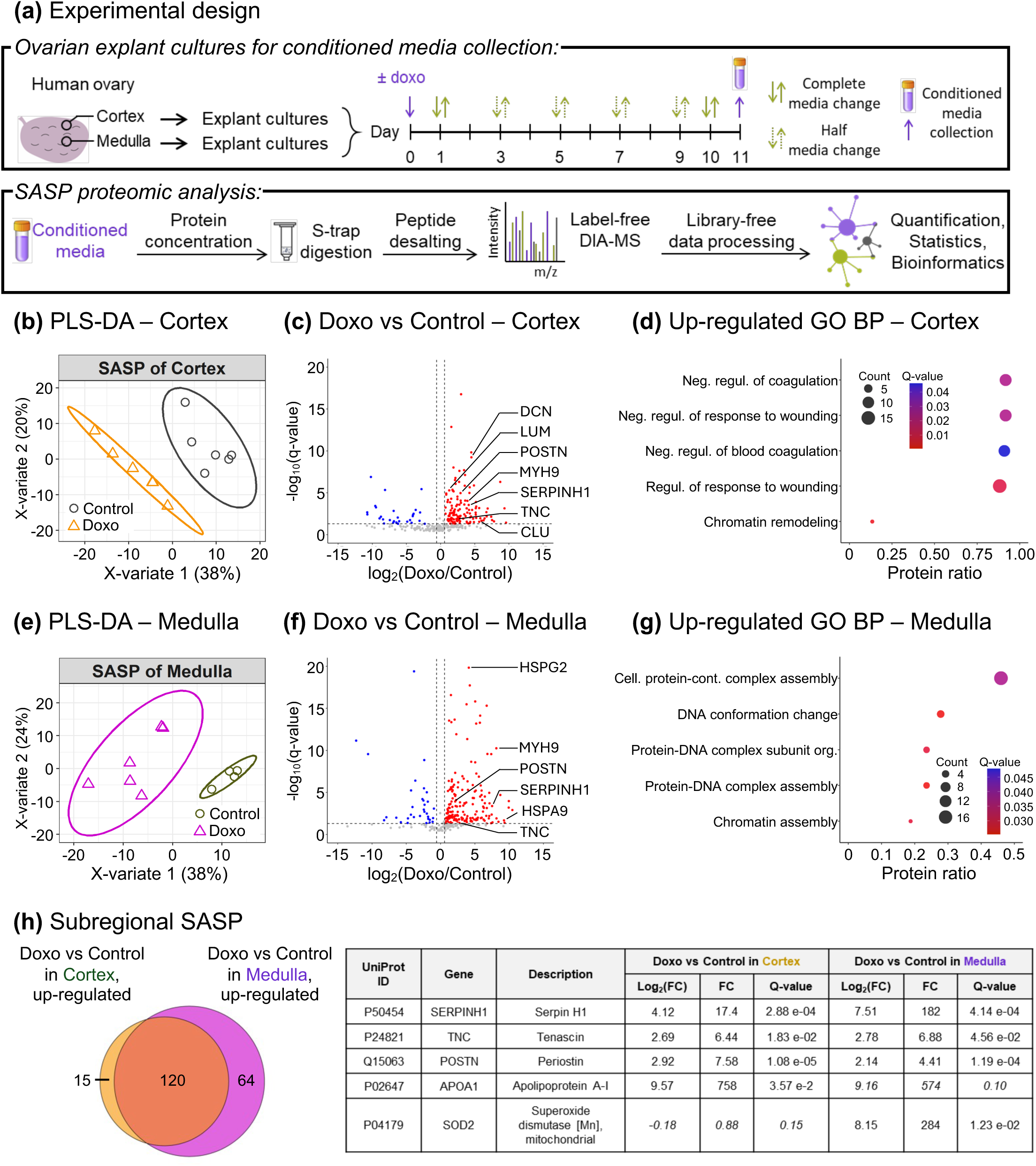
Spatial proteomic profiling of human ovarian explants identified SASP factors upon senescence induction. (a) Human ovarian cortex and medulla explants were cultured and treated with 0.1 µg/mL doxorubicin (doxo) to induce senescence or DMSO as control. After 10 days, tissues were thoroughly washed with serum-free basal media and transferred to a clean plate with pre-equilibrated serum-free basal media and inserts. The conditioned media were collected after 24 hours for proteomic analysis (Cortex: N = 6 for doxo and control, respectively, Medulla: N = 6 for doxo, N = 4 for control). Proteins were concentrated with centrifuge filters, digested using S-trap, and proteolytic peptides were desalted. Peptides were analyzed on an Orbitrap Eclipse Tribrid mass spectrometer (Thermo Fisher Scientific) operated in data-independent acquisition (DIA). DIA data were processed using directDIA (Biognosys) to identify SASP factors and enriched biological processes in the human ovarian (b-d) cortex and (e-g) medulla. (b,e) Supervised clustering using partial least squares-discriminant analysis (PLS-DA) performed with all 314 quantified protein groups (with ≥ 2 unique peptides) revealed treatment-based grouping. (c,f) The volcano plots highlight the 164 significantly altered protein groups in the cortex and 217 ones in the medulla for the ‘Doxo vs Control’ comparison (q-value < 0.05 and absolute log2(fold-change) > 0.58). The blue dots represent the down-regulated protein groups and the red dots the up-regulated protein groups. The plot y-axis is zoomed and five proteins on (c) and four proteins on (f) with q-value < 3.30e-24 are not displayed. (d,g) An over-representation analysis was performed using ConsensusPathDB. The top5 Gene Ontology (GO) Biological Processes (BP) that are up-regulated in ‘Doxo vs Control’ are displayed (q- value < 0.05 and term_level ≥ 4). (h) The Venn diagram shows the overlap between the cortex SASP and the medulla SASP, and five SASP factors are listed in the table (italic means non- significantly altered protein in the ovarian subregion).

Supervised clustering using partial least squares-discriminant analysis (PLS-DA) performed with all 314 quantified protein groups (with ≥ 2 unique peptides) revealed treatment- based grouping where the SASP from control explants (“Control”) clustered separately from SASP from doxorubicin-treated explants (“Doxo”) for both cortex and medulla (Figure 5b & 5e). In the cortex, 164 protein groups (135 upregulated and 29 downregulated) and in the medulla, 217 protein groups (184 upregulated and 33 downregulated) were significantly altered with doxorubicin exposure as compared to controls, including Basement membrane-specific heparan sulfate proteoglycan core protein (HSPG2), Myosin-9 (MYH9), and Periostin (POSTN) (q-value <0.05 and absolute log2(fold-change) > 0.58) as highlighted in the volcano plots (Figure 5c & 5f). These included factors such as Decorin (DCN), Lumican (LUM), Clusterin (CLU) in the cortex, HSPG2, mitochondrial Stress-70 protein (HSPA9) in the medulla, and POSTN, MYH9, Serpin H1 (SERPINH1), and Tenascin (TNC) in both cortex and medulla (Figure 5c & 5f). An over-representation analysis performed using ConsensusPathDB revealed Gene Ontology (GO) Biological Processes (BP) upregulated in the cortex including the regulation of coagulation and response to wounding, and in the medulla including protein-DNA complexes and chromatin assembly (q-value <0.05 and term level ≥ 4) (Figures 5d & 5g). Between the 135 proteins upregulated in the cortex and 184 proteins upregulated in the medulla, 120 proteins overlapped across both compartments with doxorubicin exposure (Figure 5h). The top five upregulated proteins were Serpin H1, Tenascin, Periostin, Apolipoprotein A1 (APOA1), and Superoxide dismutase 2 (SOD2), several of which are known to be involved in cellular senescence and aging. For example, Periostin and Tenascin-C were revealed to be crucial players in ovarian aging (Dipali et al., 2023). Additionally, Serpin H1 and Periostin appear to be biomarker candidates for chronic inflammation-associated cancers (Bons et al., 2023; González-González & Alonso, 2018). Several of these proteins are part of the established core SASP, including Periostin, Tenascin, SerpinH1, HSPG2, and COL12A1 (N. Basisty et al., 2020). However, our study also revealed potential novel SASP factors specific to the postmenopausal human ovary, including Hemopexin, Complement C5, small leucine-rich proteins (SLRPs), Nidogens, and Fibulin-1, among others (Supplemental Data 8).

### An integrated omics senotype of doxorubicin-induced cellular senescence in the human ovary

To obtain an integrated doxorubicin-induced ovarian senescence senotype in the human postmenopausal ovary, we identified the upregulated transcripts and secreted proteins that overlapped between the tissue transcriptome and conditioned media proteome for both cortex and medulla explants (doxorubicin vs control). A total of 26 unique markers overlapped between the transcriptome and the SASP, and several of these genes/proteins have been implicated in cellular senescence, aging, and/or ovarian cancer (Table 1, Figure 6a). The senotype included regulated transcripts and secreted proteins involved in extracellular matrix (ECM) organization and remodeling (LUM, LRP1, LAMA4), reactive oxygen species (SOD2, PRDX1, P4HB), metabolic pathway regulators (NAMPT, NT5E, PKM, ALDOA), cytoskeletal and actin-binding proteins (MYH9, ACTB, GSN), the protein disulfide isomerase (PDI) family of ER proteins (PDIA3, PDIA4, PDIA6), heat shock proteins (HSP90AB1, HSP90B1, HSPA5, HSPA8, HSPB1), and others (TGM2, UBC, VCP, ANXA2, CFI) (Table 1). Unbiased IPA revealed enrichment of genes in pathways including “unfolded protein response,” “protein ubiquitination,” and “cellular response to heat stress,” and this is particularly interesting given the known involvement of these pathways in cellular senescence (Abbadie & Pluquet, 2020; Hamazaki & Murata, 2024) (Figure 6b).

**Figure 6.**
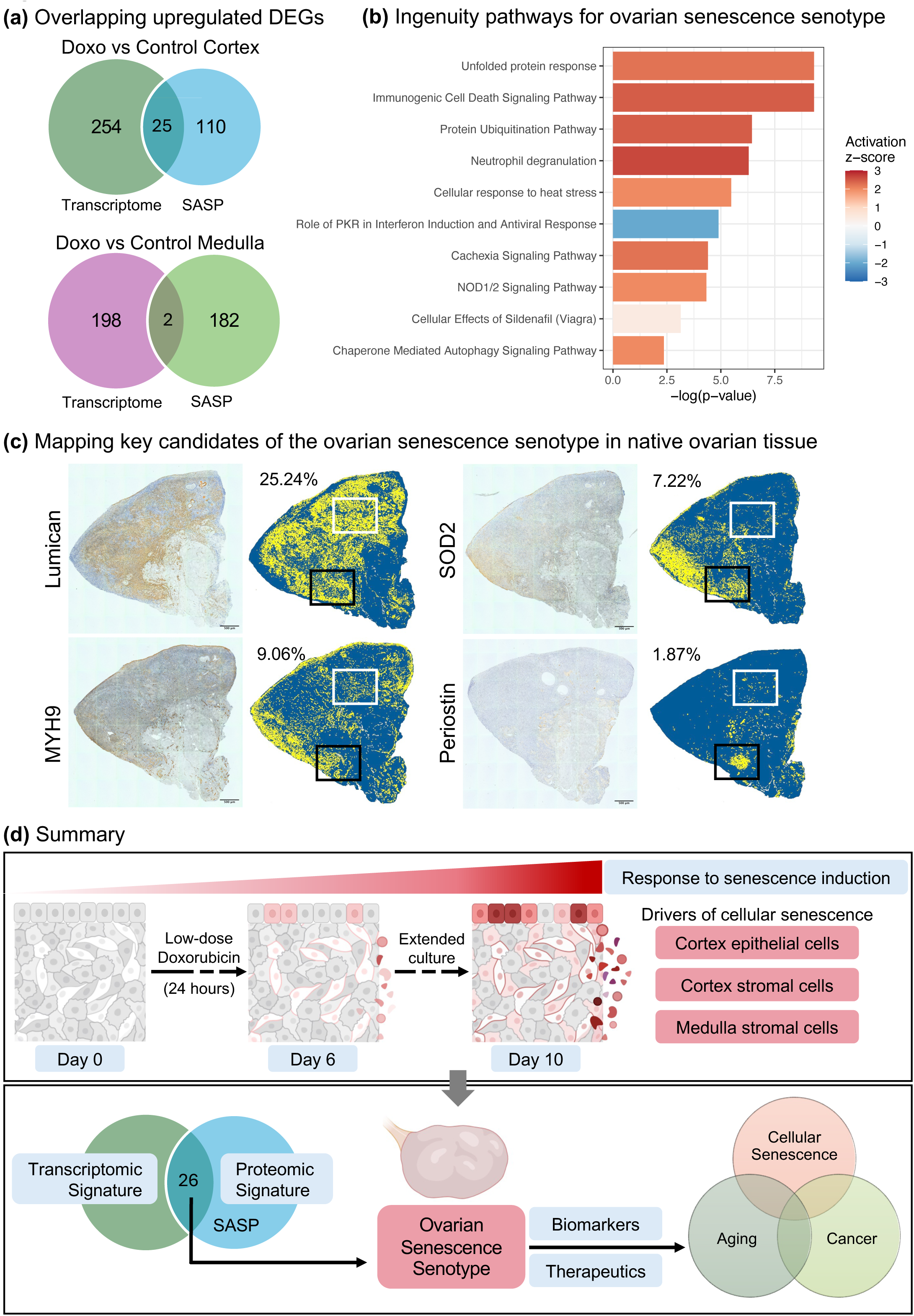
Key candidates of doxorubicin-induced cellular senescence and mapping on native postmenopausal ovarian tissue. (a) Venn diagrams showing the overlap of upregulated DEGs identified through transcriptomic and proteomic analysis in cortex (top) and in medulla (bottom). (b) Ingenuity Canonical Pathway (IPA) analysis of 26 DEGs defines the signature of doxorubicin-induced cellular senescence in the human postmenopausal ovary. (c) Mapping Lumican, SOD2, MYH9 and SASP factor Periostin on native ovarian tissue. Images on the left show colorimetric IHC scans (scale bar = 200 µm). Images on the right show digitally labelled images with yellow for positive staining and blue for negative staining. Values depict % of positive staining relative to tissue area. White and black outlined boxes represent the overlap Lumican, SOD2, MYH9, and Periostin in cortex and medulla, respectively. (d) A schematic overview of our study capturing the cellular senescence signature of doxorubicin-induced human ovarian explant model and its potential implications.

**Table 1:**
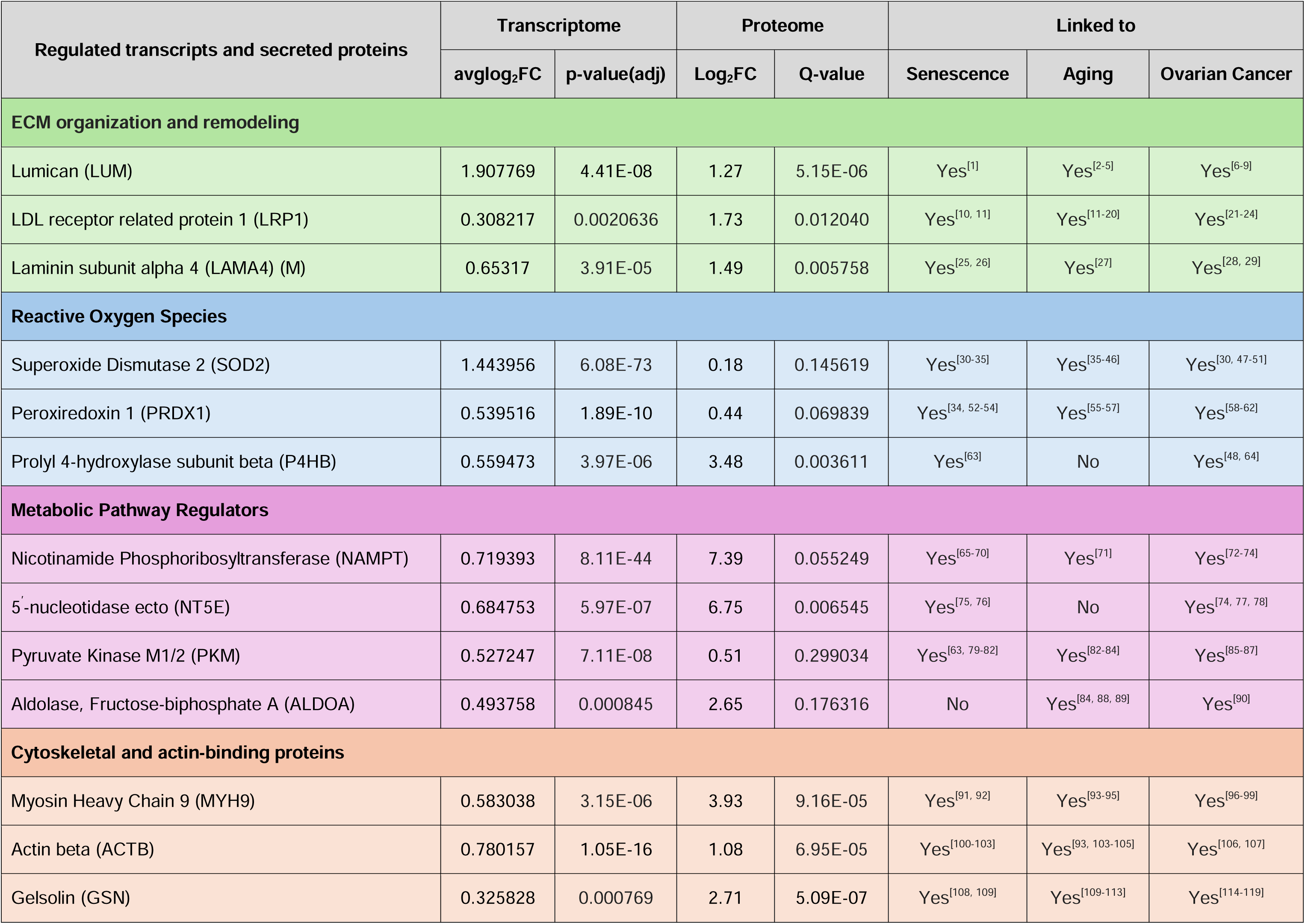
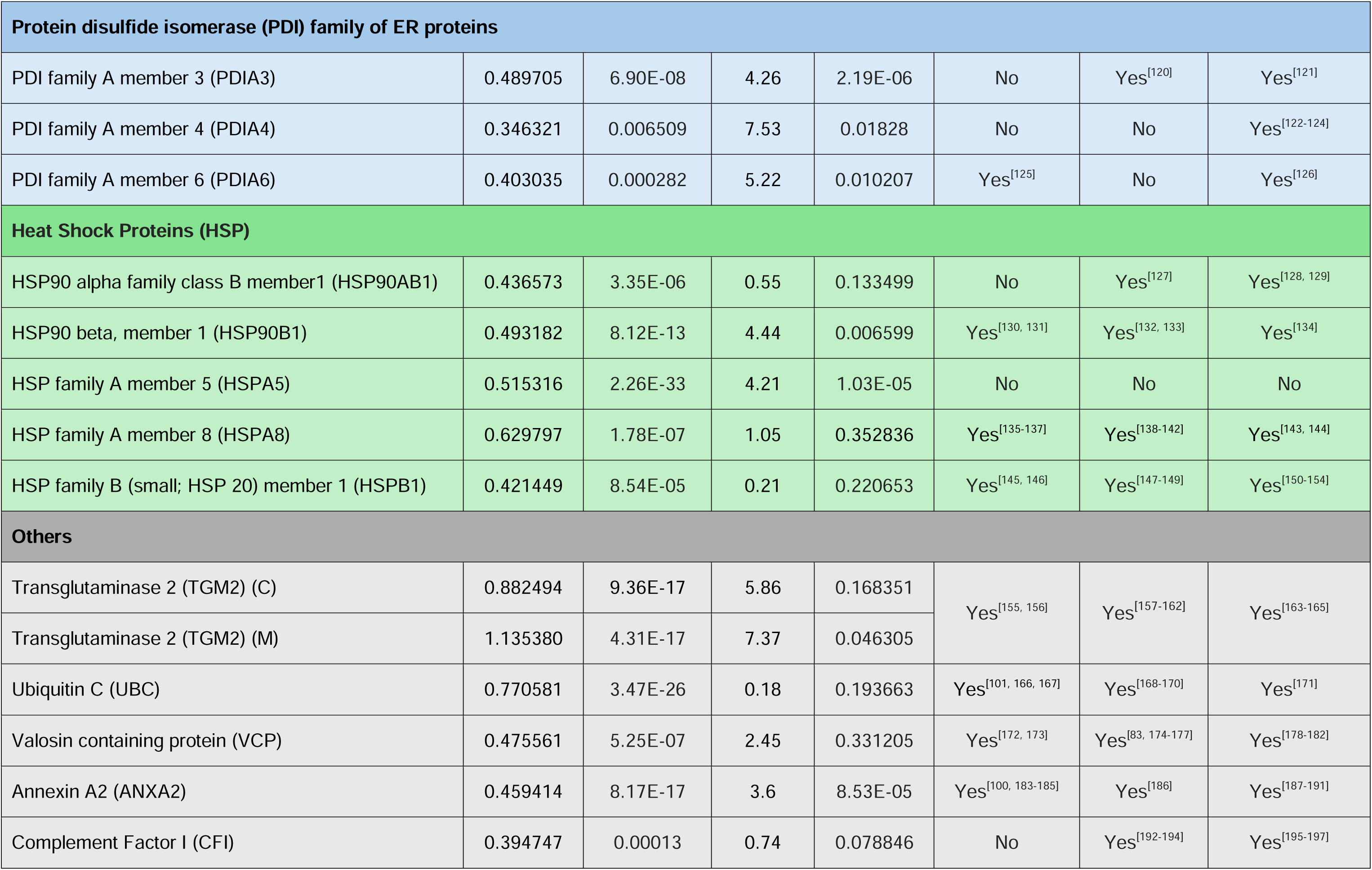
List of 26 unique DEGs and their associated proteins overlapping between the tissue transcriptome and conditioned media proteome for 10-day cultures. For transcriptome, the values represent the average log fold change (avglog2FC) and the adjusted p-value (p- value(adj)), and for proteome, the values represent the log fold change (Log2FC) and Q-values for doxorubicin vs control explants and conditioned media respectively. All DEGs are upregulated in the cortex, except LAMA4 (M) and TGM2 (M). TGM2 (C) is upregulated in the cortex.

To determine the physiologic relevance of these findings, we mapped a subset of these proteins back onto native postmenopausal ovarian tissue (Figure 6c & S8). Among these were Lumican (LUM), Superoxide dismutase (SOD2), and non-muscle Myosin IIA all of which showed the highest overlap between the transcriptome and the proteome. Additionally, we mapped SASP factor Periostin (PSTN) which was among the top 5 upregulated proteins in the cortex and medulla SASP along with SOD2 (Figure 5). All these markers were expressed in a subset of cells throughout the ovarian cortex and medulla (Figure 6c & S8). Lumican was most widely expressed (∼10-25% of tissue area), whereas periostin showed the least expression (∼0.2-2% of total tissue area) (Figure 6c, S1e, S1h, & S8a). SOD2 was expressed in both cortex and medulla (3-5% of total tissue area) with particularly strong expression in the ovarian surface epithelium (Figure 6c, S1f, and S8b). MYH9 was highly expressed in the vessel walls across both ovarian compartments (4-7% of total tissue area) (Figure 6c, S1g, & S8c). It is important to note that not all cells that stain positive for individual markers are likely to be senescent, but it may instead be regions of overlapping expression that define senescent cells (e.g. white box in cortex and black box in medulla, Figure 6c). Future studies to map and quantify these factors across a broader aging series of human ovaries will provide novel insight into how the senotype changes across age, will aid in identifying senescence biomarkers specific to the ovary, and delineate the potential role of senescent cells in ovarian aging.

## Discussion

Interrogating senescent cells in the ovary has historically been challenging due to the lack of a universal marker of cellular senescence as well as the phenotypic and causal diversity of these cells. Here we developed an induced model of cellular senescence using a doxorubicin-treated ovarian explant culture system to reveal a senescence signature in the postmenopausal human ovary using transcriptomics and proteomics approaches along with established histological markers of senescence. We established viable cultures of human ovarian explants as confirmed by histology, lack of apoptosis, and continued glucose consumption. The cortex and medulla exhibit distinct responses to senescence induction, with epithelial cells in the cortex and stromal cells in both the cortex and medulla being largely responsible. We also identified 26 unique regulated transcripts and secreted proteins overlapping between the ovarian transcriptome and secreted proteome, representing a robust integrated senotype of doxorubicin-induced senescence in the postmenopausal human ovary. Mapping key candidates from the induced senotype back to native human ovaries confirmed expression in a subset of cells, suggesting distinct roles in ovarian aging and cellular senescence (Figure 6d).

To the best of our knowledge, this is the first study investigating cellular senescence in the human postmenopausal ovary using an explant culture model (Suryadevara et al., 2024). Cellular senescence has traditionally been studied in cell culture, beginning with Hayflick and Moore’s discovery of replicative senescence in serially passaged human diploid cells (Hayflick & Moorhead, 1961). While senescent cells in culture display distinguishing and well characterized hallmarks, cell culture models fail to incorporate critical parameters like cellular heterogeneity, and intra-tissue and intercellular communication that may greatly affect the senescence phenotype (Suryadevara et al., 2024). Though some organoid models incorporate heterogeneity by using multiple cell types, they still lack the complexity and physiological relevance of tissue and its microenvironment (Adamus et al., 2014; Dos Santos et al., 2015; Lehmann et al., 2020; Uchida et al., 2019). Thus, while cell culture is a simpler system to study cellular senescence, removing cells from their complex 3D-native state in tissue and studying them in 2D controlled microenvironment greatly diminishes their translational value (Baker et al., 2011; Folgueras et al., 2018). Our model provides an intact, heterogeneous, and physiologically relevant tissue system to study cellular senescence in the human ovary.

Several chemotherapeutic agents including doxorubicin have been studied for their potential to induce cellular senescence alone, or in combination in various tissues (Alimirah et al., 2020; Altieri et al., 2016; Du et al., 2022; Kitada et al., 2019; W. Li et al., 2014; Marques et al., 2020; Uruski et al., 2021). Although there is limited characterization of histological senescence markers in human ovarian tissue (Wu et al., 2024), systemic doxorubicin in mice led increased SA-β-gal positivity, expression of p21^CIP1^ and p16^INK4a^, and levels of SASP genes (Gao et al., 2023). The observed trend towards increased expression of p21^CIP1^ and p16^INK4a^ both day 6 and day 10 of culture, along with detected SA-β-Gal activity, suggested senescence activity in our explant culture system. The aged, postmenopausal ovarian tissue is expected to have inherent levels of established senescence, and since culture itself is a senescence inducer, our measured response to doxorubicin exposure are key findings. By separately processing cortex and medulla, we were able to appreciate distinct responses to senescence induction via histology, which was further validated by transcriptomics, revealing a higher senescence score and number of DEGs in the cortex following doxorubicin treatment. In specific, the stromal cells exhibit a high senescence score, and this is consistent with a recent study demonstrating expression of senescence-related genes in the stromal cells of the postmenopausal human ovary (E. Lengyel et al., 2022). Epithelial cells also had a high senescence score, which is particularly interesting as the ovarian surface epithelium (OSE) is the primary epithelial cell type present in the postmenopausal human ovary, given the absence of granulosa or theca cells.

Given that senescence is a multifaceted cellular state involving alterations in both nuclear and secretory phenotypes, one of the key strengths of our study is the identification of a robust transcriptomic and proteomic senotype of induced senescence. In addition to novel SASP factors like Fibulin-1, Hemopexin, and SLRPs, both, transcriptomics and proteomics identified several factors that have been previously reported as senescence-associated genes including TIMP1, SERPINE1, TNC, POSTN, among others (Figure 4c, 4d) (N. Basisty et al., 2020; Coppé et al., 2010; E. Lengyel et al., 2022). These factors ranged from ECM proteins to mitochondrial and ER proteins, and several of them are known to be expressed in the human ovary and play a role in cellular senescence in other cells and tissue types (Jiang et al., 2022; Kedem et al., 2022; Li et al., 2021; Tatone et al., 2006; Velarde et al., 2012). The upregulation of ECM factors is interesting given the evidence supporting interplay between ECM factors and senescent cells and that aberrant ECM can contribute to the senescence phenotype in chronic fibrotic diseases (Blokland et al., 2020; Levi et al., 2020). Periostin, a top upregulated SASP factor is an ECM remodeling matricellular protein that increases with age in murine ovaries and is known to be implicated in ovarian cancer recurrence, along with its implication in other cancers associated with chronic inflammation (Bons et al., 2023; Dipali et al., 2023; González-González & Alonso, 2018; Huang et al., 2023; Tilman et al., 2007).

In the human ovary, the age-associated increase in fibroinflammation and tissue stiffness occurs in part due to alterations and remodeling in the ECM (Amargant et al., 2020; Dipali et al., 2023; Umehara et al., 2022). Additionally, mitochondrial dysfunction plays a crucial role in age-associated fertility decline due to oxidative stress (Grindler & Moley, 2013). All these findings are consistent with the presence of senescent cells in the ovary and may also explain the upregulation of ECM, mitochondrial, and other stress-related proteins in our integrated senotype. We identified the UPR as one of the enriched pathways in our merged signature, initiated by the accumulation of misfolded proteins. Although UPR inducers are known to trigger key hallmarks of senescence, the relationship between UPR and senescence appears to be cell-state and cell-type dependent (Abbadie & Pluquet, 2020; Druelle et al., 2016). While growing evidence highlights the role of UPR in ovarian development and function (Huang et al., 2017), its connection to senescence and ovarian aging requires further investigation. In parallel, the enrichment of other protein degradation systems, including ubiquitination and lysosomal pathway (such as chaperone-mediated autophagy), suggests a disruption in proteostasis that may contribute to the senescence signature observed in the postmenopausal human ovary. A limitation of our study is the small sample size and participant variability inherent when performing studies with healthy human tissues. Nevertheless, our multi-pronged and integrated omics approach revealed a defined ovarian senotype which can now be extended to a larger sample size to provide a broader view of cellular senescence across cell types and age.

Senotherapeutics is an emerging field focused on the development of therapeutic agents targeting cellular senescence to treat and potentially reverse aging and age-related disorders. There are conflicting results on the efficacy of senolytic treatments on reducing specific senotypes and improving ovarian aging in the mouse, which underscores the need for further investigation of cellular senescence in physiologically relevant models (Gao et al., 2023; Garcia et al., 2024; Wu et al., 2024) Our ovarian explant model holds significant potential for evaluating the effects of senotherapeutics in human ovaries (Figure 6d). Profiling cellular senescence and identifying ovary-specific senotypes will open new targets for developing these drugs. This will aid in diminishing the burden of senescent cells in the ovary and potentially extend reproductive longevity.

## Acknowledgements

We acknowledge Shriya Shah and Asma Giornazi for their help with ovarian tissue acquisition, and Kelsey Andersen, Yuna Lee, and Melania Anton for their help with ovarian tissue cultures and IHC. We also acknowledge Bernice Frederick, HTL(ASCP), Senior Research Technologist at the Pathology Core Facility of the Robert H. Lurie Comprehensive Cancer Center at Northwestern University for her help with tissue cryosections, and Brendy Fountaine and Dr. Chuck Frevert from the Histology and Imaging Core, University of Washington for their expertise and help with Visiopharm labeling and quantification for IHC.

## Authors’ contributions

F.E.D, BSch, and S.M. conceived the project, supervised the study, and provided resources. P.R.D, M.A.W, BSoy, J.B, F.W, H.A, J.P.R, T.A, N.M, T.T, E.S, D.F, J.C, M.E.G.P, S.M, BSch, and F.E.D were involved in experimental design and execution as well as data analysis, validation, and visualization. P.R.D, M.A.W, and BSoy, wrote the original draft of the manuscript that was reviewed and edited by S.M., BSch, and F.E.D. All authors have reviewed and agreed to the published version of the manuscript.

## Data availability statement

All data needed to evaluate the conclusions in the paper are present in the paper and/or the Supplementary Materials. Accession number/link for the snRNA- seq data is pending. Raw data and complete MS data sets have been uploaded to the Mass Spectrometry Interactive Virtual Environment (Mass IVE) repository, developed by the Center for Computational Mass Spectrometry at the University of California San Diego, and can be downloaded using the following FTP link: ftp://MSV000095932@massive.ucsd.edu or via the MassIVE webpage: https://massive.ucsd.edu/ProteoSAFe/dataset.jsp?task=f12d75c8d2d441d18d666f4f59098238 (MassIVE ID number: MSV000095932; ProteomeXchange ID: PXD056100).

[Note to the reviewers: To access the data repository MassIVE (UCSD) for MS data, please use: Username: MSV000095932_reviewer; Password: winter].

---

**Supplemental Figure 1.**
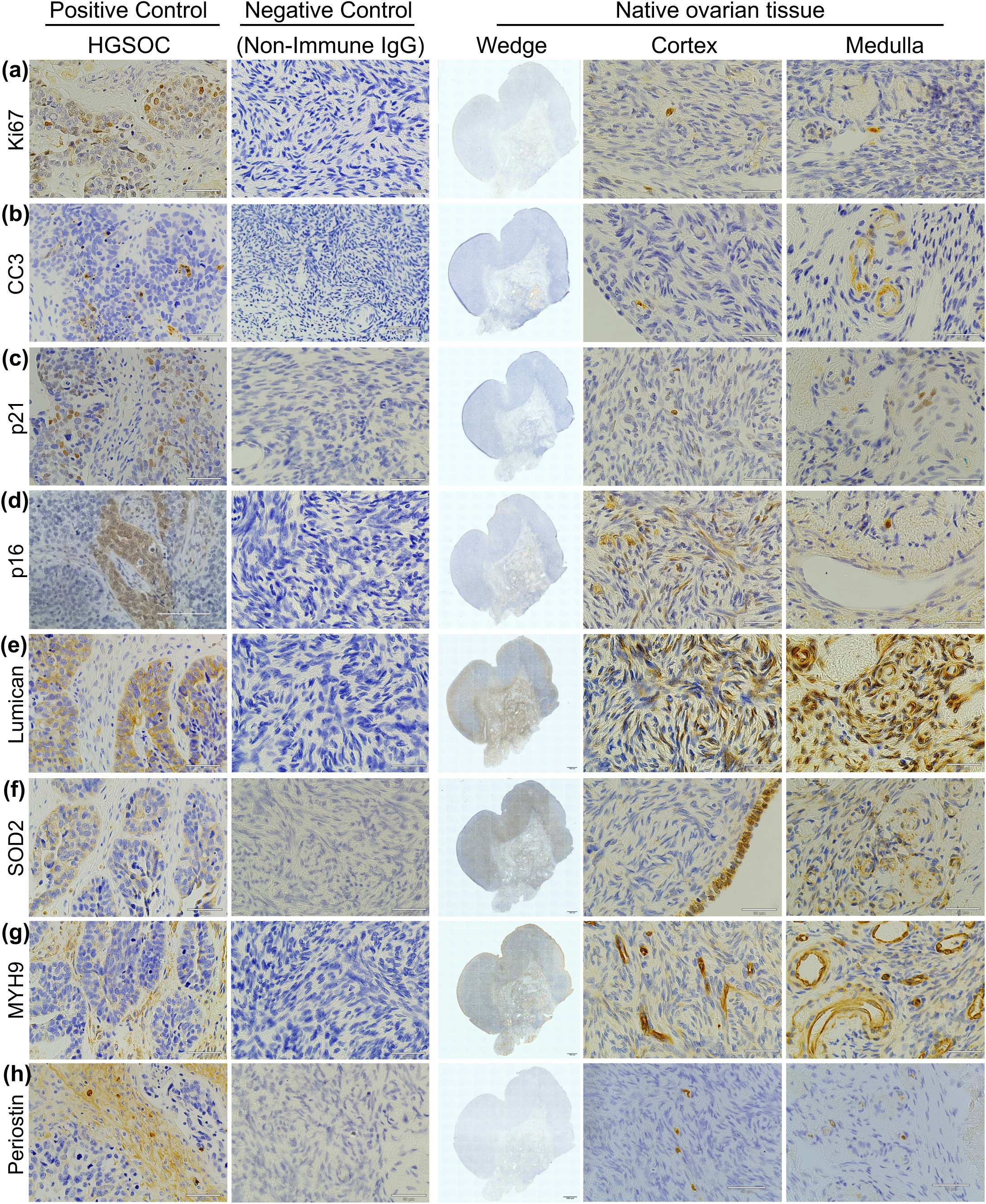
Optimization of IHC markers in native postmenopausal ovarian tissue. All IHC markers were optimized in native postmenopausal ovarian tissue, along with relevant controls including High Grade Serous Ovarian Carcinoma (HGSOC; positive control) and negative controls (with non-immune IgG). Ki67 (a), CC3 (b), and p21 (c) show very low to negative expression in the native postmenopausal ovarian tissue. p16 is expressed in discrete clusters in the cortex (d ‘Cortex’) and some expression in the medulla (d ‘Medulla’). Lumican (e) showed expression in both the cortex and medulla. SOD2 (f) showed a particularly strong expression in the ovarian surface epithelium (f ‘Cortex’). MYH9 (g) was expressed across both compartments particularly in vessel walls. Periostin (h) showed minimal expression in native tissue. Representative images from 73-yrs-old participant.

**Supplemental Figure 2.**
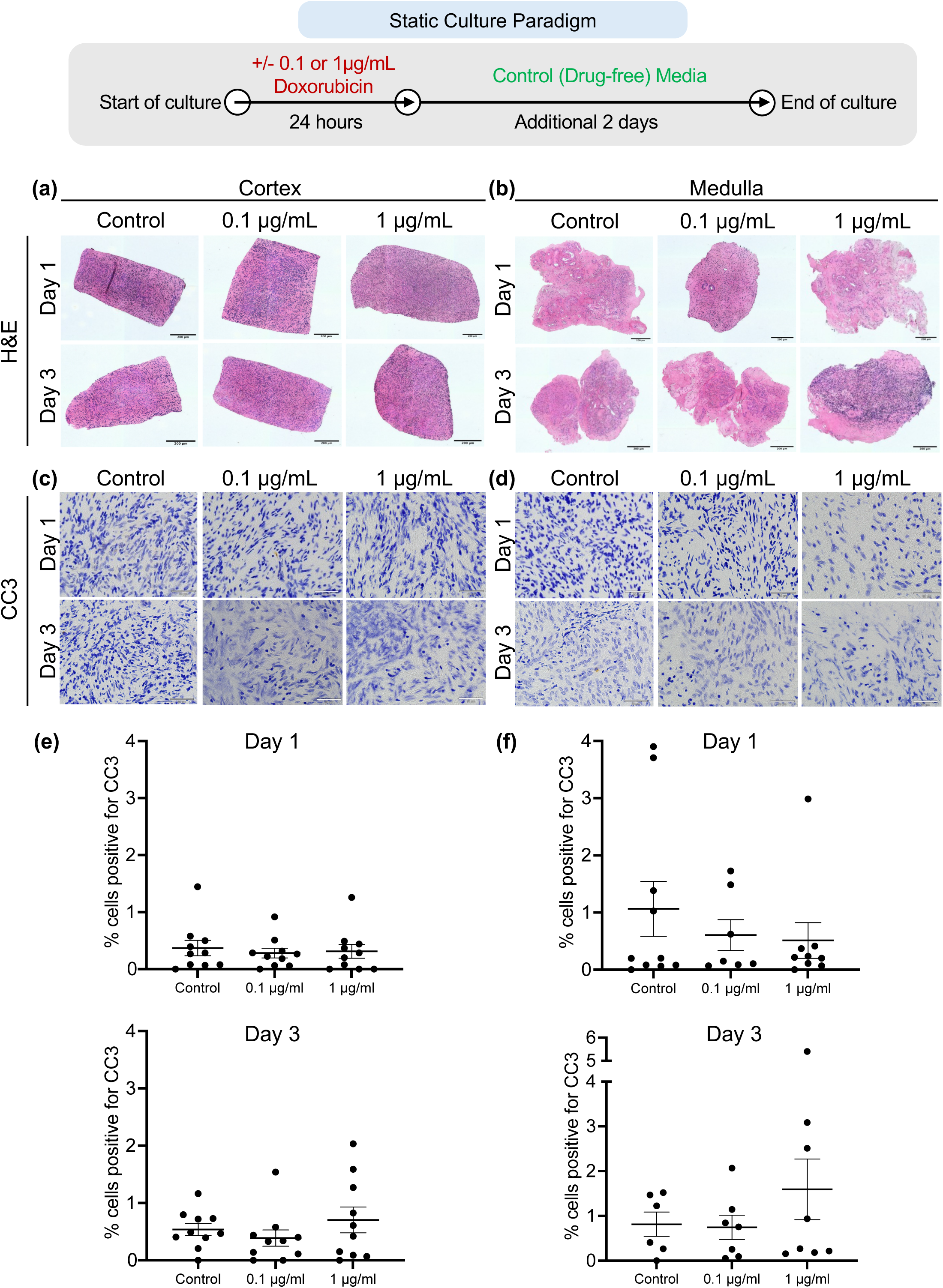
Doxorubicin dose-response and explant viability in 3-day cultures. H&E-stained sections of human ovarian cortex (a) and medulla (b) explants cultured for 2 days after 24 hours doxorubicin exposure (0, 0.1, 1µg/mL). IHC for cleaved caspase-3 (CC3) in human ovarian cortex (c,e) and medulla (d,f) explants on day 1 (Day 1) and day 3 (Day 3) of culture showed low levels of cellular apoptosis (p-value >0.05). Statistical significance was determined using an unpaired t-test and ANOVA and p-values <0.05 were considered statistically significant (N=4 participants, ages 57, 63, 68, and 70 years old). Scale bars correspond to 200 µm.

**Supplemental Figure 3.**
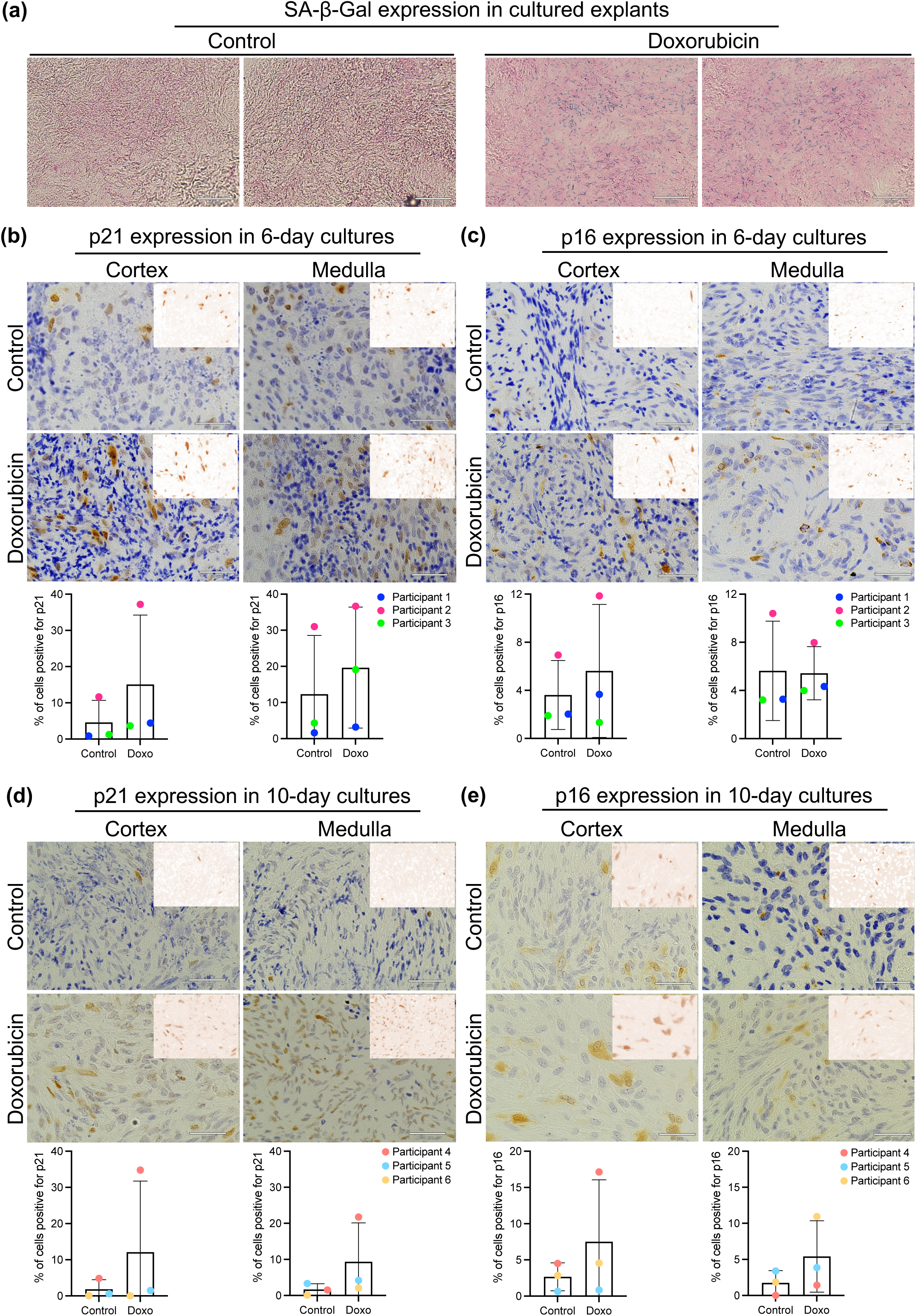
Canonical markers of cellular senescence SA-β-Gal, p21^CIP1^, and p16^INK4a^ trend towards increased expression with doxorubicin treatment. (a) SA-β-Gal staining in control vs doxorubicin-treated cultured explants. Blue color is indicative of positive SA-β-Gal staining (N=1 participant 69-yrs-old). p21^CIP1^ and p16^INK4a^ showed a trend towards increased expression with doxorubicin exposure in 6-day (2b-c; p-value Cortex =0.41 (p21) 0.61 (p16) and Medulla= 0.61 (p21) 0.94 (p16)) and 10-day (2d-e; p-value Cortex= 0.42 (p21) 0.43 (p16) and Medulla 0.29 (p21) 0.33 (p16)) cultures as compared to control. The delta in p21^CIP1^ and p16^INK4a^ expression between treated and control explants was higher after 10 days than 6 days of culture. The expression also showed variability between participants. Statistical significance was determined using an unpaired t-test and p-values <0.05 were considered statistically significant (N=3 participants for 6-day cultures; 61, 64, and 65 years old. N=3 participants for 10-day cultures; 50, 53, 62 years old). Scale bars correspond to 200 µm.

**Supplemental Figure 4.**
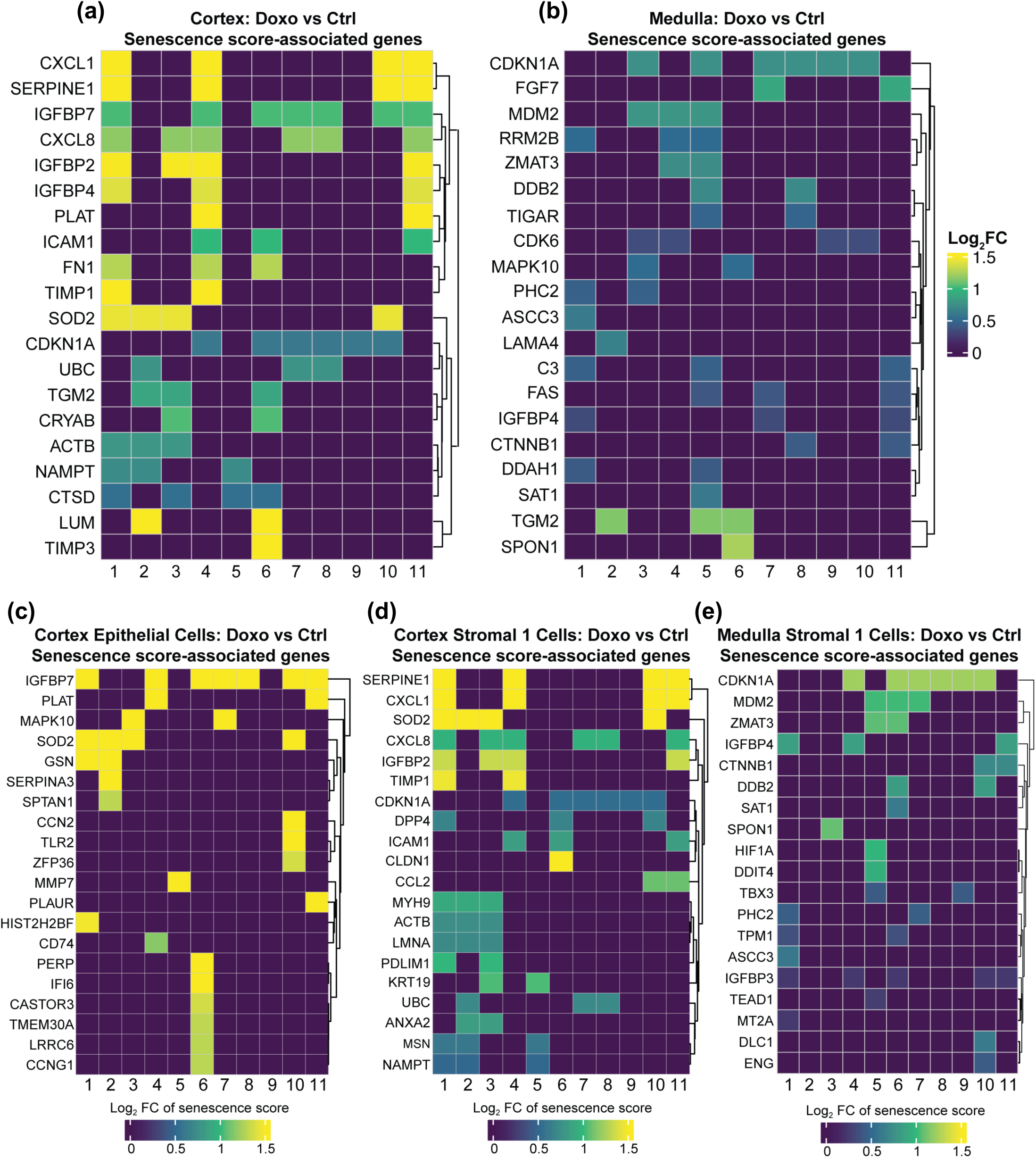
Senescent genes contributing to senescence score for ovary regions and specific cell types. The cellular senescence-associated genes that drive the day 10 senescence score in the ovarian cortex (a), medulla (b), cortex epithelial cells (c), cortex stromal 1 cells (d), and medulla Stroma 1 cells (e). The Log2FC is the comparison result of gene expression between the doxo-treated and control samples.

**Supplemental Figure 5.**
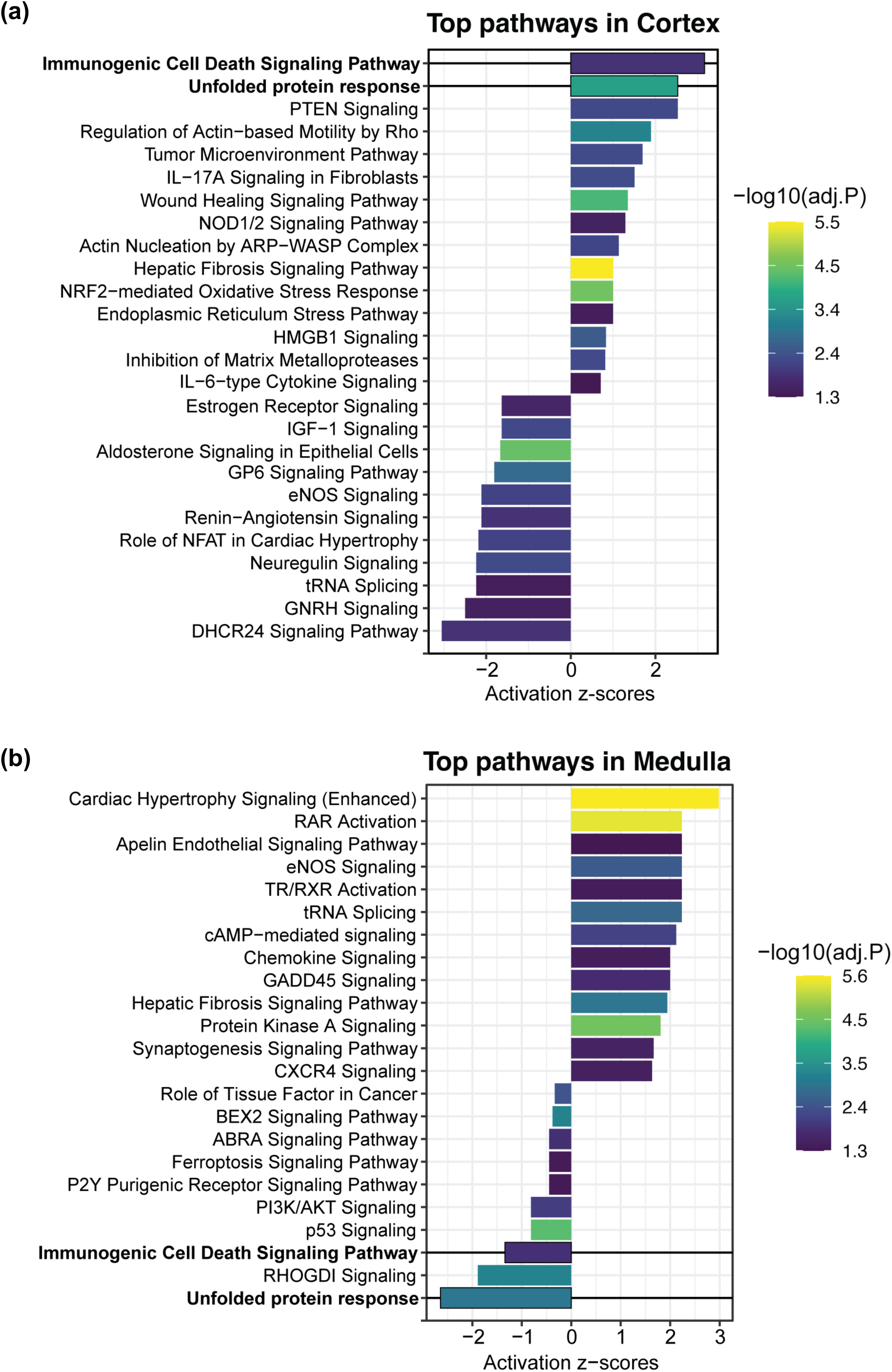
Top pathways activated and inactivated in cortex and medulla. Ingenuity Pathway Analysis (IPA, Qiagen) was used to discover the top enriched pathways in (a) Cortex, and (b) Medulla. DEGs with adjusted p-values <0.05 and Log2FC >0.2 were incorporated into the IPA canonical pathway analysis.

**Supplemental Figure 6.**
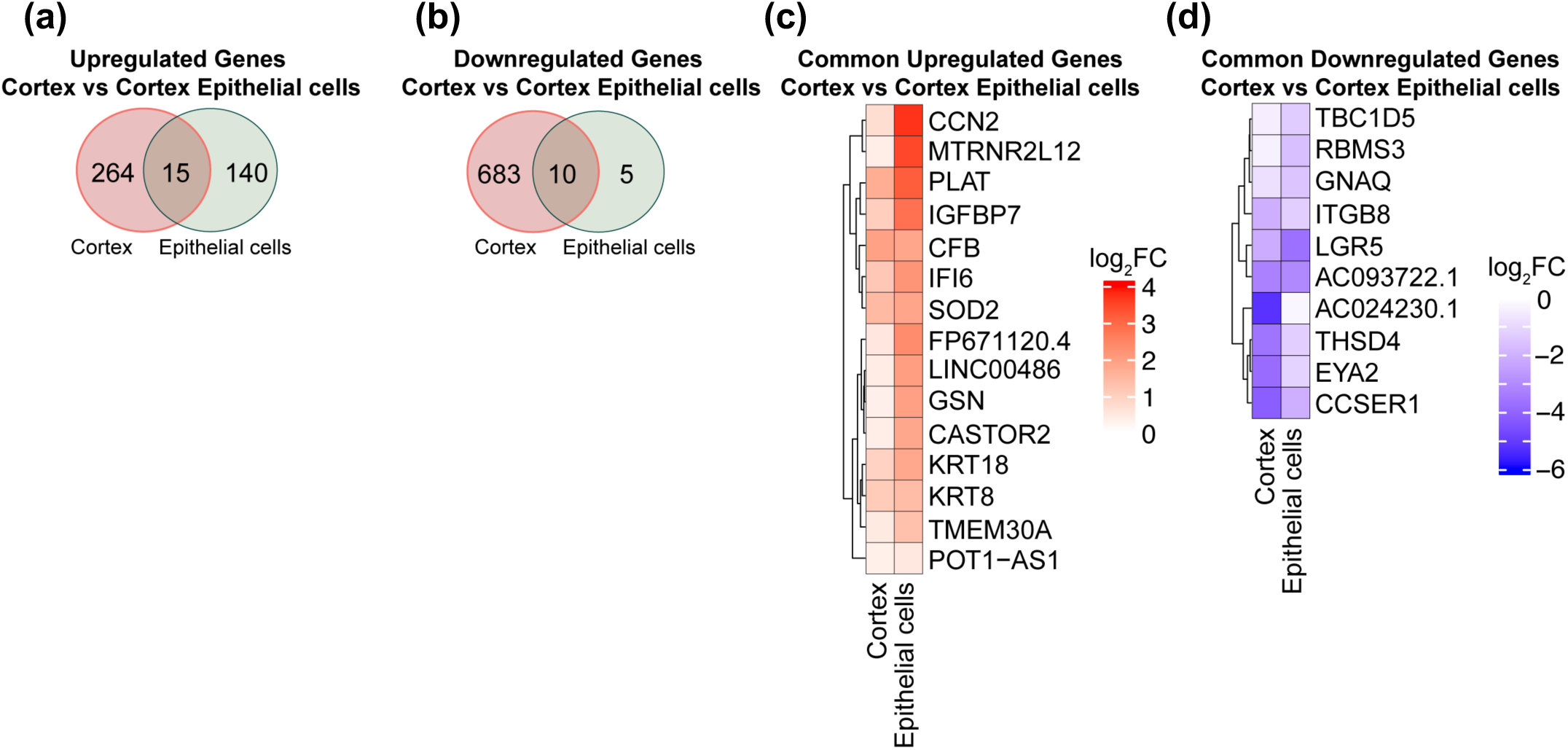
Common genes between cortex and cortex epithelial cells. (a) Venn diagram showing the overlap of upregulated DEGs between day 10 cortex and cortex epithelial cells. (b) Venn diagram showing the overlap of downregulated DEGs between day 10 cortex and cortex epithelial cells. (c-d) Heatmaps depicting the Log2FC of the 15 shared upregulated DEGs, and the 10 downregulated DEGs in the cortex and cortex epithelial cells.

**Supplemental Figure 7.**
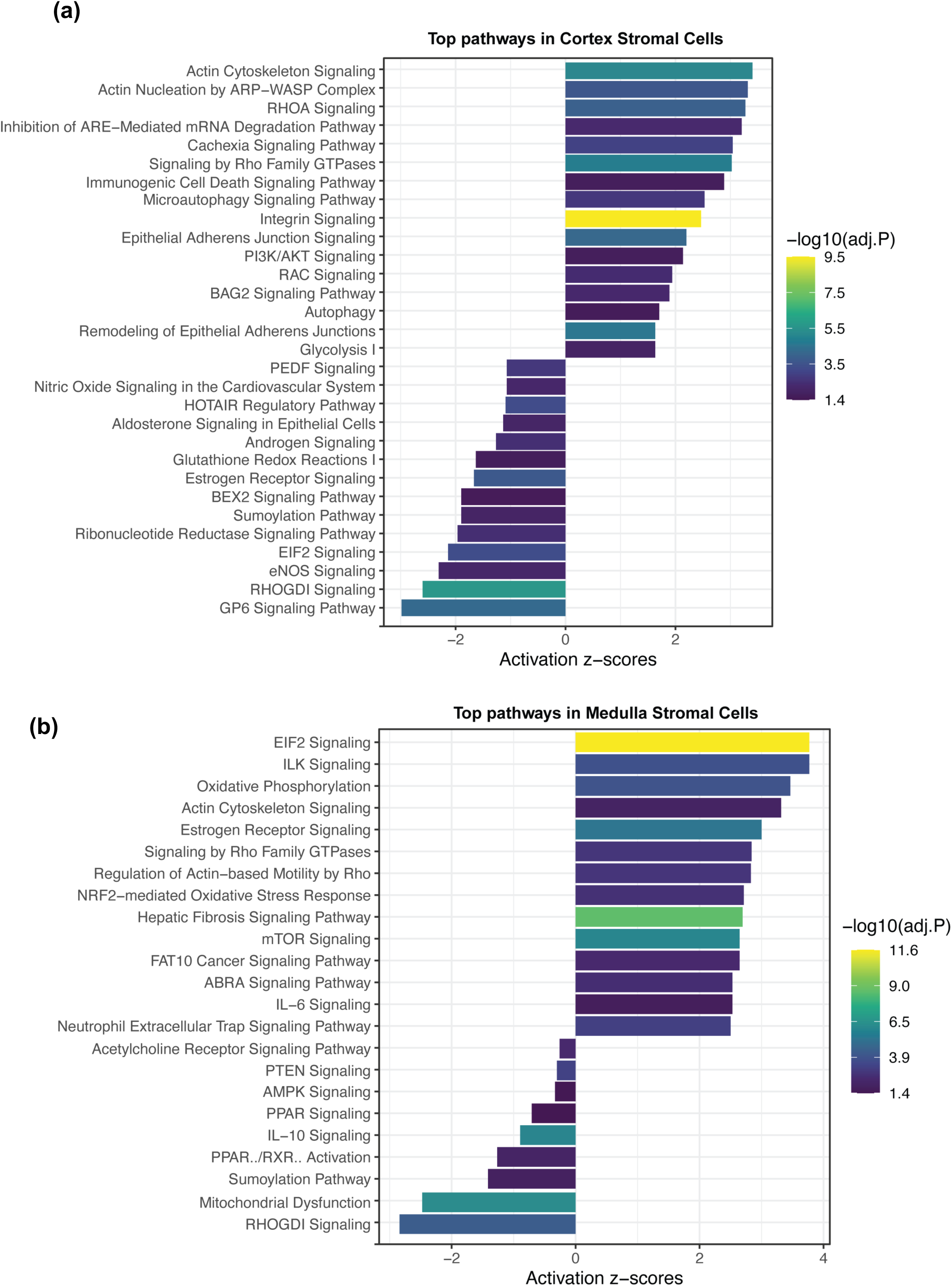
Top pathways activated and inactivated in cortex and medulla stromal cells. Ingenuity Pathway Analysis (IPA, Qiagen) was used to discover the top enriched pathways in (a) Cortex stromal cells, and (b) Medulla stromal cells. DEGs with adjusted p-values <0.05 and Log2FC >0.2 were incorporated into the IPA canonical pathway analysis.

**Supplemental Figure 8:**
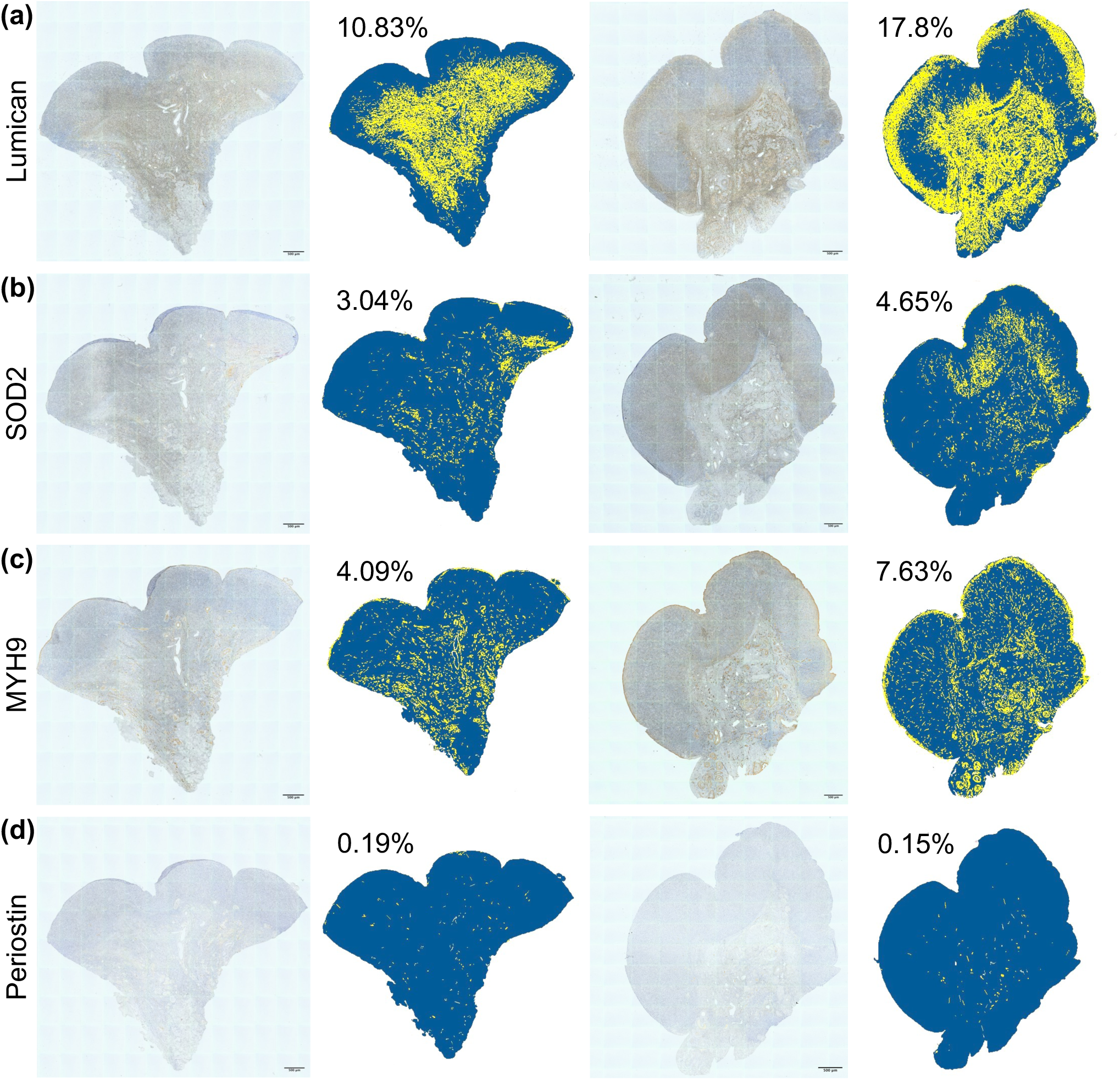
Mapping key senotype candidates in native ovarian tissue. Among the 26 proteins overlapping between the transcriptome and proteome, we mapped Lumican (a), SOD2 (b), MYH9 (c) and SASP factor Periostin (d) in native postmenopausal ovarian tissue. Images on the left show colorimetric IHC scans (scale bar = 200µm). Images on the right show digitally labelled images with yellow for positive staining and blue for negative staining. Values depict % of positive staining relative to tissue area. Representative images from 56-yrs-old and 73-yrs-old participant).

