## Supplemental Table 1 for "A comprehensive multi-omics signature of doxorubicin-induced cellular senescence in the postmenopausal human ovary"

**Supplemental Table 1**: Participant demographics and experimental information

| **#** | **Age** | **Diagnosis** | **BMI** | **Race** | **Ethnicity** | **Smoking (Y/N)** | **Gravida** | **Para** | **Experiment** | **Endpoints** |
| --- | --- | --- | --- | --- | --- | --- | --- | --- | --- | --- |
| 1 | 61 | Post-menopausal bleeding; H/O of polyps | 24.24 | White | Non-Hispanic/Latino | N | 3 | 3 | 6-day culture with or w/o 0.01µg/ml doxorubicin  for 24 hours | Morphology, Viability, Senescence markers (p16, p21), sNucSeq |
| 2 | 64 | Uterovaginal Prolapse | 23.4 | White | Non-Hispanic/Latino | N | 2 | 2 | 6-day culture with or w/o 0.01µg/ml doxorubicin  for 24 hours | Morphology, Viability, Senescence markers (p16, p21), sNucSeq |
| 3 | 65 | Cervical Dysplasia | 28.18 | African American | Non-Hispanic/Latino | N | 3 | 3 | 6-day culture with or w/o 0.01µg/ml doxorubicin  for 24 hours | Morphology, Viability, Senescence markers (p16, p21), sNucSeq |
| 4 | 50 | Post-menopausal bleeding | 34.36 | White | Non-Hispanic/Latino | N | 0 | 0 | 10-day culture with or w/o 0.01µg/ml doxorubicin  for 24 hours | Morphology, Viability,  Senescence markers (p16, p21),  SOD2, LUM, MYH9, Periostin, sNucSeq |
| 5 | 53 | Endometrial Cancer | 27.57 | White | Non-Hispanic/Latino | N | 0 | 0 | 10-day culture with or w/o 0.01µg/ml doxorubicin  for 24 hours | Morphology, Viability,  Senescence markers (p16, p21),  SOD2, LUM, MYH9, Periostin |
| 6 | 62 | Cervical Intraepithelial Neoplasia Grade 2 | 28.21 | White | Non-Hispanic/Latino | N | 2 | 2 | 10-day culture with or w/o 0.01µg/ml doxorubicin  for 24 hours | Morphology, Viability,  Senescence markers (p16, p21),  SOD2, LUM, MYH9, Periostin |
| 7 | 54 | Endometrial Hyperplasia | 19.83 | Not listed | Non-Hispanic/Latino | N | 0 | 0 | 10-day culture with or w/o 0.01µg/ml doxorubicin  for 24 hours | sNucSeq |
| 8 | 65 | Cervical Dysplasia | 22.98 | White | Non-Hispanic/Latino | N | 4 | 2 | 10-day culture with or w/o 0.01µg/ml doxorubicin  for 24 hours | sNucSeq |
| 9 | 58 | Endometrial Cancer | 30.31 | White | Non-Hispanic/Latino | N | 0 | 0 | 10-day culture with or w/o 0.01µg/ml doxorubicin  for 24 hours + Last 24 hours in basal media | Proteomic analysis of conditioned media for SASP factors |
| 10 | 56 | Uterovaginal Prolapse (Stage 3) | 22.8 | White | Non-Hispanic/Latino | N | 6 | 3 | 3-day culture with doxorubicin  dose response | Morphology, Viability (CC3) |
| 11 | 63 | Uterovaginal Prolapse (Stage 3); Urge urinary incontinence | 28.9 | African American | Non-Hispanic/Latino | Y (Former) | 3 | 3 | 3-day culture with doxorubicin  dose response | Morphology, Viability (CC3) |
| 12 | 70 | Pelvic organ prolapse (Stage 3) | 27.9 | White | Non-Hispanic/Latino | N | 3 | 2 | 3-day culture with doxorubicin  dose response | Morphology, Viability (CC3) |
| 13 | 68 | Pelvic organ prolapse (Stage 3) | 24.45 | White | Non-Hispanic/Latino | N | 5 | 4 | 3-day culture with doxorubicin  dose response | Morphology, Viability (CC3) |
| 14 | 69 | Pelvic organ prolapse (Stage 3), Stress urinary incontinence | 22.69 | White | Non-Hispanic/Latino | N | 2 | 2 | 8-day culture with or w/o 0.01µg/ml doxorubicin  for 24 hours | Senescence-associated beta galactosidase assay |
