## Supplemental Table 2 for "A comprehensive multi-omics signature of doxorubicin-induced cellular senescence in the postmenopausal human ovary"

**Supplemental Table 2:** Primary Antibody information

| **Marker** | **Source** | **Dilution** | **Concentration** | **Identifier** |
| --- | --- | --- | --- | --- |
| CC3 | Cell Signaling Technology  Danvers, MA, USA | 1:100 | 0.55 µg/mL | Asp175 (D3E9) #9597 |
| Ki67 | Dako  Santa Clara, CA, USA | 1:100 | 0.46 µg/mL | M724029-2 |
| p21^CIP1^ | Dako  Santa Clara, CA, USA | 1:100 | 2.29 µg/mL | M720229-02 |
| p16^INK4a^ | Enzo Life Sciences  Farmingdale, NY, USA | 1:300 | 2.67 µg/mL | ENZ-ABS377-0100 |
| SOD2 | Thermo Fisher Scientific, Waltham, MA, USA | 1:1000 | 1 µg/mL | MA1-106 |
| MYH9 | Abcam  Cambridge, MA, USA | 1:500 | 0.19 µg/mL | ab138498 |
| Lumican | Abcam  Cambridge, MA, USA | 1:1000 | 1 µg/mL | ab168348 |
| Periostin | Abcam  Cambridge, MA, USA | 1:500 | 1 µg/mL | ab215199 |
